# Portrait of a genus: the genetic diversity of *Zea*

**DOI:** 10.1101/2021.04.07.438828

**Authors:** Lu Chen, Jingyun Luo, Minliang Jin, Ning Yang, Xiangguo Liu, Yong Peng, Wenqiang Li, Liu Qing, Yuejia Yin, Xinnan Ye, Jiali Yan, Qinghua Zhang, Xiaoting Zhang, Songtao Gui, Shenshen Wu, Yuebin Wang, Yun Luo, Chengling Jiang, Min Deng, Min Jin, Liumei Jian, Yanhui Yu, Maolin Zhang, Xiaohong Yang, Matthew B. Hufford, Alisdair R. Fernie, Marilyn L. Warburton, Jeffrey Ross-Ibarra, Jianbing Yan

## Abstract

Maize is a globally valuable commodity and one of the most extensively studied genetic model organisms. However, we know surprisingly little about the extent and potential utility of the genetic variation found in the wild relatives of maize. Here, we characterize a high-density genomic variation map from ~700 genomes encompassing maize and all wild taxa of the genus *Zea*, identifying over 65 million single nucleotide polymorphisms (SNPs), 8 million Insertion/Deletion (InDel) polymorphisms, and over one thousand novel inversions. The variation map reveals evidence of selection within taxa displaying novel adaptations such as perenniality and regrowth. We focus in detail on evidence of convergent adaptation in highland teosinte and temperate maize. This study not only indicates the key role of hormone related pathways in highland adaptation and flowering time related pathways in high latitude adaptation, but also identifies significant overlap in the genes underlying adaptations to both environments. To show how this data can identify useful genetic variants, we generated and characterized novel mutant alleles for two flowering time candidate genes. This work provides the most extensive sampling to date of the genetic diversity inherent in the genus *Zea*, resolving questions on evolution and identifying adaptive variants for direct use in modern breeding.

## Introduction

Global crop production is currently unable to meet the anticipated demands of a growing human population (**Ray DK et al. 2013; Bailey-Serres J et al. 2019**). Further exacerbating the problem in many areas, crops are being impacted by climate change (**Lesk C et al. 2016**), and projected shifts in temperature and precipitation will lead to further declines in productivity for many major crops (**Challinor AJ et al. 2014**). New varieties displaying both higher yield and the ability to adapt to diverse environments are thus urgently needed to increase crop productivity under changing climate scenarios (**Tigchelaar M et al. 2018; Li Q and Yan J. 2020**).

Maize (*Zea mays* ssp. *mays*) is one of the world’s most widely grown crops, with an annual global production of over 1.1 billion tons in 2018 (FAOSTAT, 2020). It was domesticated from the wild grass *Zea mays* ssp. *parviglumis* (hereafter *parviglumis*) approximately 9,000 years ago in the southwest of Mexico (**Matsuoka Y et al. 2002; Piperno DR et al. 2009**). Population genetic analyses largely agree that maize underwent a substantial population bottleneck during domestication (**Eyre-Walker A et al. 1998; Tenaillon MI et al. 2004; Wright SI et al. 2005; Beissinger TM et al. 2016**), reducing the genetic diversity available for adaptation. Although maize rapidly spread from its center of domestication across a wide range of environments, successful adaptation required hundreds or thousands of years (**Swarts K et al. 2017**). As global populations increase and climate change accelerates, unprecedented maize yield losses are projected to become commonplace in most maize-producing regions (**Tigchelaar M et al. 2018; Zampieri M et al. 2019**). To facilitate adaptation to these new challenges, breeders will need to maximize use of the genetic diversity at their disposal, looking beyond modern elite lines to traditional cultivated varieties and locally adapted wild relatives (**Zhang H et al. 2018**).

The wild congeners of maize, collectively called teosinte, are annual and perennial grasses native to Mexico and Central America (**Figure 1a**). They are adapted to a diverse range of environments, from hot, humid, tropical regions of Central America to cold, dry, high elevations of the Mexican Central Plateau (**Hufford MB et al. 2012a; Sánchez González JJ et al. 2018**). Teosinte exhibit novel biotic and abiotic adaptations absent in modern maize (**Mammadov J et al. 2018**), providing a wealth of genetic diversity that could be utilized in modern breeding. A recent example is a large-effect allele for leaf angle identified in teosinte that was lost during maize domestication (**Tian J et al. 2019**): CRISPR-Cas9 editing of maize to mimic the teosinte allele resulted in a ~20% yield increase in modern hybrids grown at high density. Other studies have also used genetic mapping to capitalize on teosinte alleles for nutrition (**Karn A et al. 2017; Li K et al. 2019**), extreme environments (**Mano Y et al. 2012; Mano Y and Omori F. 2013**), and disease resistance (**Lange ESD et al. 2015; Lennon JR et al. 2015; Lennon JR et al. 2017**). Population genetic evidence suggests that diverse alleles from the teosinte *Zea mays* ssp. *mexicana* (hereafter *mexicana*) played an important role in allowing maize to adapt to arid highland conditions (**Hufford MB et al. 2013**).

**Figure 1.**
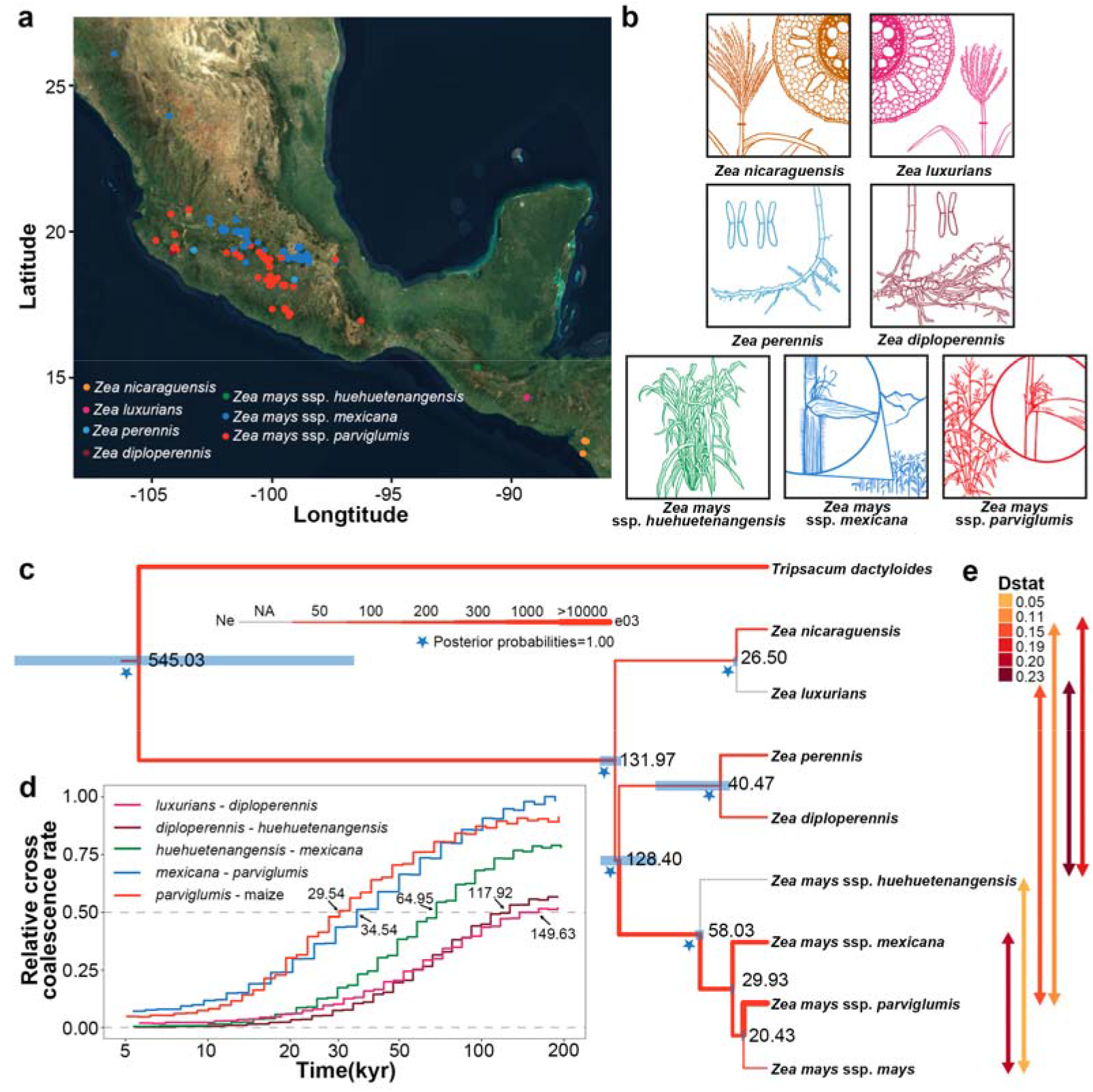
Phylogeny of *Zea* genus. **a**, Geographical distribution of collected teosinte, populations were identified and colored based on morphology. **b**, Morphological characteristics of teosinte (Credit to Dr. Andi Kur). *nicaraguensis* and *luxurians* are distinguished from other teosinte based on aerenchyma in their stems which aerate roots during submergence, *nicaraguensis* has a more robust tassel than *luxurians*. *perennis* is a recent autotetraploid of *diploperennis*, both of these perennial taxa are distinguished from other teosinte based on their rhizomatous root systems. The Mexican annual teosinte *parviglumis* and *mexicana* are distinguished from each other based on the presence of macro-hairs and pigment along their stems, two traits that are linked to highland adaptation. **c**, Divergence times estimated from the multispecies coalescent (MSC) model. Blue bars indicate the 95% highest posterior density (HPD) intervals. The star indicates nodes with posterior probability of 1. Edge widths reflect estimates of effective population size (**Supplementary Table 4**). **d**, MSMC2 cross-population results among teosinte species. Curves between *luxurians* and *huehuetenangensis* were computed using two phased haplotypes, and other groups were computed using four phased haplotypes. **e**, Introgression among taxa. Arrows indicate the taxa involved, and arrow color shows the value of Patterson’s D-statistic (**Supplementary Table 5**).

Despite the vast potential for teosinte to contribute to breeding and adaptation of cultivated maize, we know relatively little about the genetic diversity and history of these taxa. Estimates of the age of the genus vary wildly (**Gaut BS and Clegg MT. 1993; Hilton H and Gaut BS. 1998; White SE and Doebley JF. 1999; Ross-Ibarra J et al. 2009; Wang Q and Dooner HK. 2012**), and the phylogenetic relationship of several taxa are debated or unknown (**Doebley JF. 2005; Hufford MB et al. 2012a**). There is considerable cytological diversity in the genus, as large inversions (**Ting YC. 1965; Ting YC. 1967; Ting YC. 1976; Fang Z et al. 2012; Pyhäjärvi Y et al. 2013; Yang N et al. 2017**) and transposable element variation (**Lamb JC and Birchler JA. 2006; Tenaillon MI et al. 2011; Chia JM et al. 2012**) have been documented in some taxa. Moreover, common garden studies have demonstrated that much of the phenotypic differentiation in both teosinte and maize landraces is the result of local adaptation (**Fustier MA et al. 2019; Janzen GM et al. 2021**). Low density genotyping or pooled sequencing approaches in *parviglumis* and *mexicana* have identified a number of candidate loci related to soil, climate, and disease resistance, highlighting the importance of inversions (**Pyhäjärvi T et al. 2013; Fustier MA et al. 2017; Aguirre-Liguori JA et al. 2019**). However, for most taxa in *Zea*, their potential as sources of useful diversity in maize are poorly understood.

Here, we present a genus-wide resource of genome-scale genetic diversity in *Zea*. We resequenced 183 teosinte accessions to high depth and combined these data with sequences from 507 maize inbred lines. Our analyses reveal a detailed phylogeny and demography of the genus *Zea*, including frequent gene flow among taxa. We identify substantial novel genetic diversity for SNPs and small InDel polymorphisms, and over 1,200 inversion polymorphisms, of which 14 are larger than five megabases. We investigate selection over long time scales among all taxa and identify candidate regions associated with local adaptation in highland teosinte (*mexicana*). We also used this deep sampling of maize lines to dissect the genetics of adaptation to temperate climate and highlight convergent adaptation of highland teosinte and high-latitude maize. Finally, we demonstrate the utility of our population genomics approach to identify novel functional diversity, using genome editing to knock out two candidate flowering time genes and validate their effects in a maize background. Together, our integrated study of evolutionary history, genetic diversity and local adaptation in the genus *Zea* will facilitate the efficient use of diverse *Zea* taxa in modern maize breeding and improvement.

## Results

### The haplotype map and phylogeny of the genus *Zea*

We resequenced 183 teosinte accessions encompassing all described species and subspecific taxa in the genus *Zea* (**Figure 1a, b**), and combined these with genome resequencing data from 507 cultivated maize inbred lines representing those grown in both temperate and tropical regions (**Yang XH et al. 2011; Supplementary Table 1**). To ensure the quality of this new *Zea* haplotype map, we used a set of strict filtering conditions (**Online methods**), detecting a total of 65,717,599 SNPs (**Supplementary Table 2**), of which 79% were rare variants (MAF<0.05) (**Supplementary Figure 1**) and 8,601,262 small InDels. We validated a subset of SNPs using Sanger sequencing, with median concordance between datasets >95% and reasonable false positive (18% on average) and false negative rates (4% on average) for non-reference alleles (**Supplementary Table 3**). Based on population structure analysis, samples with greater than 60% ancestry in a single group were clustered into *parviglumis* (*n*=70), *mexicana* (*n*=81), *Zea mays* ssp. *huehuetenangensis* (*n*=1; hereafter, *huehuetenangensis*), *Zea diploperennis* (*n*=2; hereafter, *diploperennis*), *Zea perennis* (*n*=2; hereafter, *perennis*), *Zea luxurians* (*n*=1; hereafter, *luxurians*), *Zea nicaraguensis* (*n*=12; hereafter, *nicaraguensis*), and 507 diverse maize inbred lines (**Supplementary Figure 2a, b; Supplementary Table 1**). Principal component analysis of these lines was in strong concordance with population structure results (**Supplementary Figure 2c**).

We inferred phylogenetic relationships for the genus *Zea* under the multispecies coalescent model (**Figure 1c; Flouri T et al. 2018**), with maximum likelihood phylogenies (**Lee TH et al. 2014**) producing largely congruent results (**Supplementary Figure 3-4**). Notably, we estimated a very recent origin for the genus, splitting from its sister genus *Tripsacum* only ~500,000 years ago. This young age is especially striking given the pronounced differences in chromosome structure and subgenome organization resulting from the two genera’s shared polyploidy event >10M years ago (**Wang X et al. 2015**). Within the genus, our results suggest that *nicaraguensis* likely represents a subspecies of *luxurians*, with divergence times similar to those among subspecies of *Zea mays*. Our model also estimates relatively small population sizes for *diploperennis* and *nicaraguensis*, with a recent population expansion in the autopolyploid *perennis* and among the annual teosinte in *Zea mays* (**Figure 1c; Supplementary Table 4**).

The phylogeny supports an earlier analysis (**Ross-Ibarra J et al. 2009**) suggesting that divergence among *Zea mays*, *luxurians*, and *diploperennis* was nearly contemporaneous, occurring ~130,000 years ago (95% highest posterior density (HPD) interval for *luxurians* divergence from other taxa: 125,467-144,333; **Figure 1c; Supplementary Table 4**). We further estimate that *perennis* split from its diploid progenitor *diploperennis* only ~40,000 years ago (95% HPD: 31,200-95,900). Tree topologies and divergence times also support earlier analyses (**Buckler ES and Holtsford TP. 1996**) showing that *huehuetenangensis* is a subspecies of *Zea mays*, diverging from other annual subspecies ~58,000 years ago (95% HPD: 53,667-61,297), followed by the divergence of highland *mexicana* and lowland *parviglumis* ~30,000 years ago (95% HPD: 27,533-30,900). Independent estimates of the divergence time taken from rates of cross-coalescence between taxa under the Multiple Sequentially Markovian Coalescent (MSMC2; **Malaspinas AS et al. 2016**) are strikingly consistent (**Figure 1d**), but differences in the estimates between some pairs of taxa (**Supplementary Figure 5**) lead us to suggest a possible role for gene flow during species divergence (**Ross-Ibarra J et al. 2009; Warburton ML et al. 2011**). Indeed, patterns of shared derived alleles suggest a history of introgression among taxa (**Figure 1e; Supplementary Figure 6; Supplementary Table 5**). These include admixture between *parviglumis* and each of the other species of *Zea*, and between domesticated maize and both *huehuetenangensis* and *mexicana*, highlighting the important role of gene flow in crop adaptation (**Janzen GM et al. 2019**). Finally, we hypothesize that both population structure in *parviglumis* and substantial admixture with wild relatives may explain our otherwise surprising overestimate of the timing of maize domestication (~20,000 years ago) compared to the archaeological record (~9000 years ago; **Piperno DR et al. 2009**).

### Novel diversity in *Zea*

SNP data highlight the impressive genetic diversity present in teosinte. Despite the potential downward bias due to strict filtering parameters and read mapping to a maize reference, heterozygosity and nucleotide diversity are both higher in teosinte taxa than the much larger panel of maize lines, even among teosinte with limited geographic ranges (**Supplementary Table 2; Supplementary Figure 7**). Nearly half (46%; **Supplementary Table 2**) of the SNPs and 53% of the InDels identified across all taxa are taxon-specific (**Figure 2a; Supplementary Figure 8a**), and differences in the number of private SNPs correlate with estimates of the effective population size (**Supplementary Figure 8b**). The majority (~70%) of these private SNPs are relatively rare and are only found within populations at derived allele frequencies of <5%, suggesting that deeper sampling than that reported here would likely continue to uncover novel variation. Differentiation (*F*_ST_) between teosinte taxa is often lower than that found between inbred maize and teosinte (**Figure 2b; Supplementary Figure 9a**), consistent with the historical reduction of diversity that occurred during modern maize breeding (**van Heerwaarden J et al. 2012**). The annual subspecies of *mays* shows much faster decay of linkage disequilibrium than our diverse panel of maize inbreds (5-10Kb compared to ~180Kb; **Supplementary Figure 9b**), but historical recombination in *nicaraguensis* appears to be even more limited (>500Kb).

**Figure 2.**
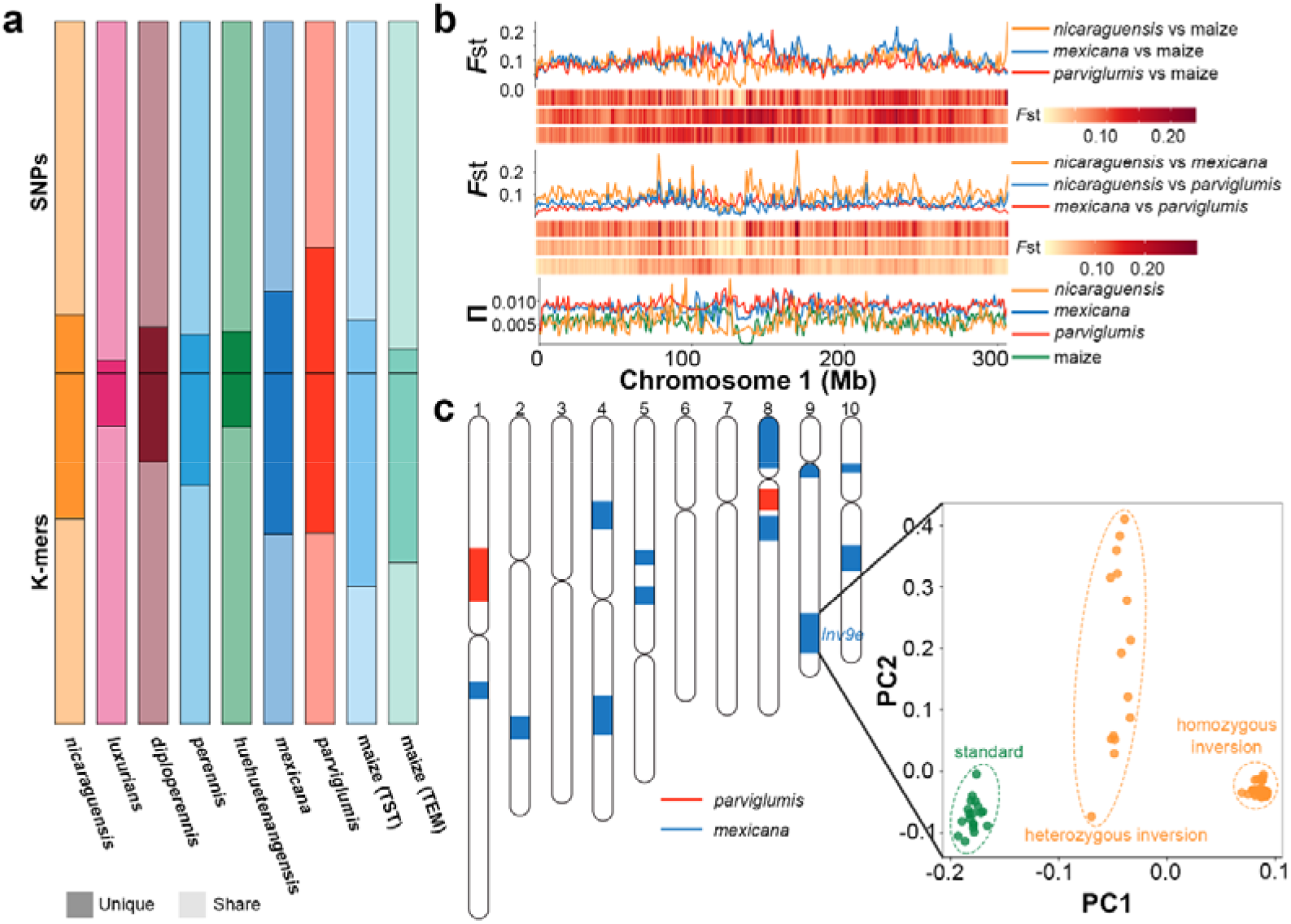
Variation in the *Zea* genus. **a**, Private SNPs and unique 31bp k-mers in *Zea* genus. Maize (TST) represents tropical maize, maize (TEM) represents temperate maize. **b**, Distribution of *F*_ST_ and nucleotide diversity along chromosome 1 in *Zea*. Heatmaps of *F*_ST_ comparisons follow the order of comparisons in the legend. **c**, Distribution of inversions across the chromosomes. Each colored segment represents an inversion, with colors referring to the population in which the inversion is most prevalent (red: *parviglumis*; deep blue: *mexicana*). Inset shows PCA of SNPs data from within *Inv9e*, clearly separating the three genotype classes (left: standard; middle: heterozygous inversion; right: homozygous inversion).

Short-read mapping approaches pose challenges in characterizing genetic diversity, including poor power to detect structural variants, difficulty with repetitive sequences, and reference bias. In order to circumvent some of these obstacles, we used a reference-free k-mer approach to characterize each taxon (**Online Methods**). Consistent with an allele frequency spectrum heavily shifted towards rare variants (**Supplementary Figure 1**), most taxa showed a substantial proportion of unique k-mers (**Figure 2a; Supplementary Table 2**). For example, we find that ~15% of k-mers in *huehuetenangensis* and *luxurians* are unique, even though both taxa are represented by only a single sample in our data. Our large sample of domesticated maize also demonstrated considerable k-mer diversity, with the majority (~60%) of k-mers observed in tropical maize not found in any of the teosinte genomes. These results not only highlight the novel genetic diversity present in teosinte but also likely point to the ongoing importance of evolutionary processes in generating and filtering diversity in traditional maize populations in Mexico (**Bellon MR et al. 2018**).

We used this data to investigate the diversity and abundance of inversion polymorphisms in *Zea*. Inversions are known to play important roles in adaptation (**Faria R et al. 2019**), and previous work has highlighted the evolutionary relevance of several large inversions in *Zea* (**Fang Z et al. 2012; Mano Y et al. 2012; Pyhäjärvi T et al. 2013; Crow T et al. 2020**). Using diagnostic changes in linkage disequilibrium across inversions (**Cáceres A and González JR. 2015**), we detected more than 1,200 large (>500Kb) genomic regions with patterns indicative of inversion polymorphism in *parviglumis* or *mexicana* (**Supplementary Table 6, 7; Supplementary Figure 10**). Of these, 14 (12 in *mexicana* and two in *parviglumis*) were larger than 5Mb (**Figure 2b; Supplementary Table 8**). Eleven of the 14 inversions are newly identified in the present study, and, in most cases, genetic diversity in these regions clearly identifies alternative haplotypes (**Figure 2b; Supplementary Figure 11**). GO enrichment analysis of genes within inversions highlights loci involved in biotic and abiotic stress response (**Supplementary Table 9; Supplementary Figure 12)**, including response to wounding (GO: 0009611) and oxidative stress (GO: 0006979). Given that many inversions found segregating at appreciable frequency are likely adaptive in some environments, these data argue strongly that improved assembly and characterization of structural variants in teosinte is a promising avenue for the discovery of genetic diversity useful for maize adaptation and breeding.

### Phylogenetic signals of selection among lineages

Their genetic, ecological, and life history diversity make teosintes an ideal model system for studying adaptation (**Hufford MB et al. 2012a**). To investigate patterns of adaptation over longer time periods, we performed a phylogenetic scan for lineagespecific rates of nonsynonymous substitutions in individual genes (**Supplementary Figure 13a; Online Methods**). We find hundreds of genes showing both positive and purifying selection in each of the taxa (**Supplementary Table 10, 11**). Consistent with a model in which *Zea* taxa show unique adaptations, genes showing evidence of positive or purifying selection are generally unique to each taxon (**Supplementary Figure 13b, c**), even among closely related taxa (**Supplementary Figure 13d-i**). For example, we identify a large number of genes showing evidence of selection on branches underlying key life history traits like waterlogging resistance in *luxurians* and *nicaraguensis* (**Supplementary Figure 13d, g**) or perenniality in *diploperennis* and *perennis* (**Supplementary Figure 13e, h**). Among the genes under purifying selection, we find suggestive enrichment in particular biological functions (**Supplementary Table 12**). For example, in the perennial taxa (*diploperennis* and *perennis*), we find enrichment in DNA recombination (GO: 0006310), telomere organization (GO: 0032200), and other biological processes related to chromosome stability. One gene under purifying selection that has been characterized previously is *ZmPHS1* (**Supplementary Table 13**), a gene involved in meiotic recombination and the homologous chromosome pairing process (**Pawlowski WP et al. 2004; Ronceret A et al. 2009**). The significance of these findings to the speciation and local adaptation of teosinte subspecies is worth further exploring.

### Local adaptation between highland and lowland teosinte

In addition to phylogenetic analysis of selection, our extensive sampling of *parviglumis* (*n*=70) and *mexicana* (*n*=81) accessions allowed investigation of more recent adaptation. We first took advantage of having the geographic location of each of our teosinte samples to perform environmental association analysis. We focus on elevation and soil characteristics (**Online Methods**), two environmental features that play important roles in local adaptation of these subspecies (**Pyhäjärvi T et al. 2013; Fustier MA et al. 2017; Aguirre-Liguori JA et al. 2019**). Across both sets of association results, we find a number of hits within *Inv9e*, a structural arrangement identified cytologically in *mexicana* (**Ting YC. 1964**) and highlighted as important for adaptation across elevation (**Pyhäjärvi T et al. 2013**).

We identified a clear peak of association with elevation within *Inv9e* (**Supplementary Figure 14a**), with the top SNP located ~8.5Kb upstream of *ZmWOX11*, a WUSCHEL-related homeobox transcription factor that key to development of plants (**Jha P et al. 2020**). Given this putative evidence for selection on regulatory sequence, we investigated patterns of gene expression and conducted further experiments. Transcripts of *ZmWOX11* were found in abundance in meristem tissues in the base of the shoot and tips of secondary roots, but at very low levels in the leaf (**Supplementary Figure 14b; Supplementary Table 14**). Transgenic lines overexpressing *ZmWOX11* exhibited significantly increased ear height and plant height in two field environments (**Supplementary Figure 14c-f**), and, consistent with observed changes in soil ions across elevation (**Supplementary Table 15; Wei S et al. 2014**), hydroponic experiments identified increased primary or secondary root length of *ZmWOX11* overexpression lines under CaCl_2_ (20mM), NaCl (100nM), PH (4) and Pi (500μm) treatments for 12 days (**Supplementary Figure 15**).

We next performed genome-wide association analysis with 29 representative traits from a rich database (**Online Methods**) of more than 200 soil properties (**Supplementary Figure 16a; Supplementary Table 15; Wei S et al. 2014**). In total, we identified 33 QTLs related to 15 soil properties (**Supplementary Figure 17; Supplementary Table 16**). *Inv9e* had previously been reported to correlate with topsoil variables (**Pyhäjärvi T et al. 2013**) and here we find it co-locates with six QTLs related to four soil traits (**Supplementary Figure 17; Supplementary Table 16**). Within the QTL, the transcription factor *Zm00001d047878* (~8.6Mb away *ZmWOX11*) is a likely candidate for a QTL for electrical conductivity and gypsum content. It is differentially expressed between *mexicana* and *parviglumis* (**Supplementary Figure 16b**) and its *Arabidopsis* ortholog, *BHLH32*, acts as a negative regulator of root hair formation in response to phosphate starvation (**Chen ZH et al. 2007**).

Adaptation to the mountains of central Mexico likely involved a host of important challenges beyond abiotic factors such as soil and elevation. To identify genes underlying highland adaptation more broadly, we scanned the genome for evidence of haplotype frequency differentiation in *mexicana*. We identify more than 3,000 regions showing evidence of such selection (**Figure 3a; Supplementary Table 17**), with a large plurality (42%, *P*<0.001 **Supplementary Figure 18**) located in one of the large inversions identified above, likely reflecting both suppressed recombination associated with the inversion and the importance of these structural variants for adaptation.

**Figure 3.**
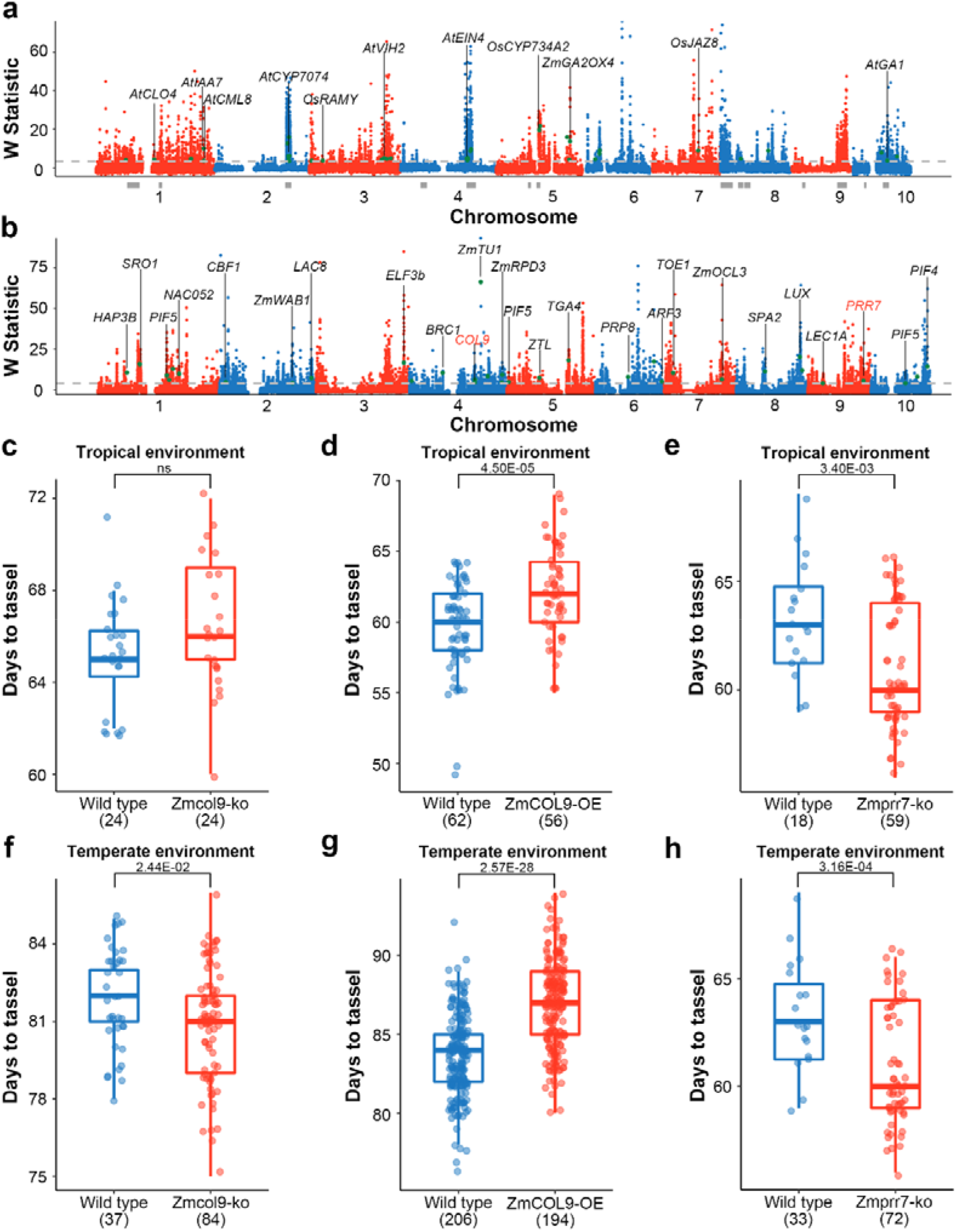
Local adaptation in teosinte and maize. **a**, Genome-wide selection signals (W statistic; smoothed XP-CLR score) between *mexicana* and *parviglumis*. The horizontal grey dashed line represents the top 5% cutoff. Genes differentially expressed under biotic and abiotic stress that have been functionally validated in maize, rice and *Arabidopsis thaliana* are labeled as green points. Grey rectangles below indicate large inversions (> 5Mb) between *parviglumis* and *mexicana*. **b**, W between temperate and tropical maize. Genes associated with flowering time and floral development in maize, rice and *A. thaliana* are marked with green points. **c-h**, Days to tassel of wild type and mutants under tropical (**c-e**; Hainan province 2019; China; E109°, N18°) and temperate (**f-g**; Jilin province 2020; China; E125°, N44°) environments; ns indicates no significant difference. Shown are comparisons of wild type and Zmcol9 CRISPR/Cas9 knockout mutants (**c**, **f**), wild type and *ZmCOL9* overexpression individuals (**d**, **g**), and wild type and Zmprr7 CRISPR/Cas9 mutants (**e**, **h**).

Plant hormones play an important role in stress response, and crosstalk between different hormones enables plants to tolerate a variety of harsh environments (**Verma V et al. 2016; Ku YS et al. 2018**). Twelve of the candidate highland adaptation genes identified based on haplotype frequency differentiation were annotated as members of plant hormone signal transduction pathways by KEGG (**Supplementary Figure 19**). Moreover, 216 candidates show evidence of differential expression under biotic or abiotic stress (**Hoopes GM et al. 2019**), and 41 of these have been functionally validated in maize, rice or *A. thaliana*, with 17 genes regulating or responding to plant hormones for stress response (**Supplementary Table 18**).

### Temperate adaptation in maize

Though maize was domesticated in the warm lowland tropics, by 4,300 BP it had spread to the higher latitudes of the southwestern United States (**Merrill WL et al. 2009**), reaching southern Canada by 2,300 BP (**Hart JB and Hetty LR. 2007**). Previous work identified evidence of convergent evolution between temperate maize and cold-adapted *Tripsacum* (**Yan L et al. 2019**), and we extended this comparison to investigate convergence between temperate maize and high elevation adaptation in teosinte. Applying the same scan for haplotype differentiation between temperate and tropical maize, we identified thousands of genes showing evidence of allele frequency differentiation across latitude (**Figure 3b; Supplementary Table 19**) and found a statistically significant overlap of genes showing evidence of selection in both highland teosinte and temperate maize (89 genes; *P* = 0.029). In addition to shared patterns of selection, we used RNA-seq from the shoot base of *parviglumis*, *mexicana* and tropical and temperate maize (**Supplementary Table 1**) to search for similar changes in gene expression. We identified 595 genes differentially expressed between *mexicana* and *parviglumis* (**Supplementary Table 20**) and 437 genes differentially expressed between tropical and temperate maize (**Supplementary Table 21**), with significant overlap between the two lists (15 shared genes; *P* = 0.001). But while genes showing evidence of selection in teosinte show greater than expected change in expression, consistent with selection targeting regulatory changes, we see no such difference in temperate maize (**Supplementary Figure 20)**.

Selection for variants that promote early flowering enabled maize to break day-length restrictions and facilitated the spread of maize across a broad geographical range (**Buckler ES et al. 2009**). Experimental data in maize and from orthologs in other taxa shows that at least 35 genes associated with high latitude adaptation were involved in flowering time pathways (**Figure 3b; Supplementary Table 22**). For example, the gene *tunicate1*, known for its dramatic impact on the pod-corn phenotype in maize (**Han JJ et al. 2012)**, shows evidence of selection in *mexicana* and the strongest selection signal in high latitude adaptation and mutants of *tunicate1* have been shown to influence flowering time of maize (**Figure 3b**).

To validate the utility of our selection scan approach, we chose two candidate genes from the photoperiod-regulated and circadian-clock-controlled flowering pathways: *ZmCOL9* (*Zm00001d051684*) and *ZmPRR7* (*Zm00001d047761*). Mutants of these two genes were obtained from a CRISPR/Cas9-based high-throughput targeted mutagenesis library (**Liu H et al. 2020**). The loss-of-function allele of *ZmCOL9* includes a 5 bp deletion/1bp insertion in the intron and a 2 bp deletion/4 bp deletion in the 3^rd^ exon (**Supplementary Figure 21a, d**) that result in premature translation termination. In a tropical environment (Hainan; China; E109°, N18°), *ZmCOL9* knockout mutants showed no difference in flowering time compared to the wild type (**Figure 3c; Supplementary Figure 21b, e**) but overexpression plants exhibit a later flowering phenotype (**Figure 3d; Supplementary Figure 22a, b)**. In contrast, when planted in a temperate environment (Jilin; China; E125°, N44°), the *ZmCOL9* knockout mutants exhibited significantly earlier flowering (**Figure 3f; Supplementary Figure 21c, f**) and the overexpression lines maintained a later flowering phenotype compared to the wild type (**Figure 3g; Supplementary Figure 22c, d**). The mutant allele of *ZmPRR7* is a 5.8-Kb deletion of a gene region which leads to the total loss of protein function. Plants harboring the mutant allele exhibit significantly earlier flowering than the wildtype in both tropical and temperate environments (**Figure 3e, h; Supplementary Figure 23**). These results confirmed the key roles for both *ZmCOL9* and *ZmPRR7* in regulating flowering time and contributing to the adaptation of modern maize.

## Discussion

The twin projections of increasing human population and decreasing available farmland amplifies the challenge breeders face in producing high crop yields without harming the environment, and this has motivated an increasing interest in crop wild relatives (**Hufford MB et al. 2012b; Aguirre-Liguori JA et al. 2017**). Here, we present a high-resolution genetic variation map which greatly expands the publicly available genetic sequence information for the genus *Zea*. All data and results of this work were integrated into the ZEAMAP database (**Gui S et al. 2020**) for easy query and retrieval.

This work provides a more complete picture of the phylogeny and demography of the genus *Zea*, including both divergence times and effective population sizes of *Zea* species. We reaffirm several aspects of the phylogeny of *Zea*, but our data identify a number of new features, including the likely subspecies status of *nicaraguensis*, the short divergence times between the perennial taxa, and the relatively young age of the genus. We caution that our divergence estimate for *Tripsacum* may be underestimated, however, and that high-quality *Tripsacum* and teosinte reference genomes will be essential to better answer this question.

Climate change and limited farmland availability represent grand challenges for modern crop improvement (**Kumar M. 2016; Xu H et al. 2016; Giannini TC et al. 2017**). After initial domestication in western Mexico, maize rapidly spread to reach its current broad geographical distribution (**Meyer RS and Purugganan MD. 2013**). Teosinte wild relatives have always been distributed over a much more limited geographical range, but this range nevertheless spans extremely diverse environmental conditions (**Hufford MB et al. 2012a**). Inversions are believed to be a major driver of adaptation and speciation (**Kirkpatrick M and Barton N. 2006; Faria R et al. 2019**). Previous studies have highlighted the importance of *Inv9e* in *mexicana* adaptation (**Pyhäjärvi T et al. 2013; Fustier MA et al. 2017; Aguirre-Liguori JA et al. 2019**), but did not identify the underlying causal genes. Here, we identified *ZmWOX11* as one of the causal genes in *Inv9e* that functions in high elevation adaptation of *mexicana*. In a genome-wide adaptive allele scan across the *Zea* genus, we also revealed lineage-specific loci related to local adaptation. By focusing on adaptation to high elevation and latitudes, we identified 1,387 and 2,057 candidate loci, respectively. In the identified adaptive regions of teosinte, we found that gene expression variation was more abundant, perhaps reflecting a loss of plasticity (**Lorant A et al. 2017**) or *cis*-regulatory variation **(Hufford MB et al. 2012c**) during domestication.

Plants are subjected to different biotic and abiotic stress during adaptation to new local environments. Many candidate highland adaptation loci identified in this study are involved in phytohormone pathways, a well-known pathway that regulates biotic and abiotic stress stimuli (**Suzuki N. 2016; Miller RN et al. 2017; Yang J et al. 2019; O’Brien AM et al. 2019**). However, it is particularly noteworthy that there are large numbers of specific highland adaptive alleles in *mexicana* which neither exist in cultivated maize nor in its direct ancestor *parviglumis*. Our functional analysis of these highland adaptation alleles clarifies the great potential in the utilization of the wild relatives of maize, which serve as a treasure of alleles for modern maize improvement. Analysis of high-latitude adaptation genes also uncovers the important role of flowering-time regulation in maize temperate adaptation and provides promising candidate loci to ameliorate yield reduction in the face of global warming. *ZmCOL9* and *ZmPRR7* were both further verified in CRISPR/Cas9 experiments, confirming the potential value of our method to identify and functionally validate genes relevant for adaptation. These data thereby suggest that combinations of population genetics analysis and next generation precision gene editing tools will greatly accelerate modern crop genetic improvements (**Fernie AR and Yan JB. 2019**). The data and discoveries presented in this study provide the foundation for the use of crop wild relative resources when breeding for climate change adaptation.

## ONLINE METHODS

### Samples and whole genome resequencing

A total of 183 teosinte accessions from CIMMYT and one *Tripsacum dactyloides* from Dr. Fajia Chen’s lab (Henan Agricultural University, China) were obtained, consisting of 90 *mexicana*, 75 *parviglumis*, 12 *nicaraguensis*, two *perennis*, two *diploperennis*, one *huehuetenangensis*, one *luxurians* and one *Tripsacum dactyloides* according to morphological classification (**Supplementary Table 1**). Young leaves from one individual of each accession were used for DNA extraction for sequencing using the Illumina HiSeq3000 platform (150-bp paired-end), conducted by BGI (Shenzhen, China). DNA sequencing data of 507 cultivated maize and 1 *Tripsacum dactyloides* were downloaded from the NCBI SRA database (PRJNA531553; SRA051245; **Supplementary Table 1**).

### Read mapping, SNP calling

Raw reads of teosinte and maize were first processed using FastQC (v0.11.3; http://www.bioinformatics.babraham.ac.uk/projects/fastqc/) and Trimmomatic (v0.33; **Bolger AM et al. 2013**) to remove poor-quality base calls and adaptors. Reads were then aligned to the B73 reference genome (v4; **Jiao YP et al. 2017**) using Bowtie2 (v2.1.0; --very-fast; **Langmead B et al. 2012**). Unique mapped reads were sorted and indexed using Picard (v1.119; http://broadinstitute.github.io/picard/). SAMtools (v1.3.1; **Li H et al. 2009**) and UnifiedGenotyper from GATK (v3.5; https://software.broadinstitute.org/gatk/) were used to estimate the variant calling file for each individual. Hard filtering of the individual SNP calls was carried out with mapping quality (MQ ≤ 20.0), and thresholds set by sequencing coverage based on minimum coverage (DP ≤ 5) and maximum coverage (DP ≥ 200). Then, variants from the 183 teosinte and 507 maize were combined by GATK CombineVariants to a single variant calling file. To confirm if unknown variants were discarded reference genotypes in individual calls, we recalled these sites and replaced them with reference genotypes if they had supported reads. Finally, sites with a missing rate higher than 75% in all samples were excluded. To validate the accuracy of SNPs called from resequencing data, 349 sites in 194 accessions were selected for Sanger sequencing (**Supplementary Table 3**).

### Population structure classification, principal component analysis and phylogenetic tree construction

We evaluated patterns of population structure using a set of SNPs filtered to remove multiallelic loci and SNPs with a minor allele frequency < 0.05 (--maf 0.05 -biallelic-only) using PLINK (v1.9; **Chang CC et al. 2015**). We then ran admixture for different values of the number of clusters (K) from 2 to 10 (--cv = 10; v1.3.0; **Alexander DH et al. 2009**). Each individual with admixture components < 0.6 was classified as ‘Teosinte (mix)’ and ‘maize (mix)’. We performed PCA analysis using this same set of SNPs with GCTA (v1.26; **Yang J et al. 2011**) recording the first 10 components (--pca 10). We annotated non-missing SNPs in teosinte and maize with SnpEff (v4.1g; http://snpeff.sourceforge.net/index.html) using the first transcript of B73 v4 genes. We then used synonymous and noncoding SNPs to construct a simple phylogenetic tree with SNPhylo using default parameters (v20140701; **Lee TH et al. 2014**) and visualized the tree with iTOL (**Letunic I and Bork P. 2019**).

### Species tree analysis

Species delimitation and species trees were inferred using BPP (model A11; v4.1.4; **Flouri T et al. 2018**). We used the following samples in BPP: 10 tropical maize, 10 *parviglumis*, 10 *mexicana*, 10 *nicaraguensis*, two *diploperennis*, two *perennis*, one *luxurians*, one *huehuetenangensis* and two *Tripsacum dactyloides* (**Supplementary Table 1**). Low-quality base calls and adaptors from raw reads of *Tripsacum dactyloides* were removed using Trimmomatic, and then aligned to the B73 v4 reference genome with Bowtie2 as described above. The consensus base was estimated from the uniquely mapped reads using ANGSD (v0.930; **Korneliussen TS et al. 2018**). Using the B73 annotation, we randomly selected 2,000 coding sequence genes to estimate the species delimitation and species tree. The prior distribution of ancestral population size (θ) and divergence time from the root (τ) followed an inverse-gamma (IG) prior with means of 0.005 IG (3 0.01)) and 0.75 (IG (3 1.5)), respectively. The consensus of A11 species trees was visualized using DensiTree (v2.2.6; **Bouckaert RR. 2010**).

### Imputation and demographic estimation

SNPs in the 183 teosinte were imputed with BEAGLE (v4.0; **Browning SR and Browning BL. 2007**). After imputation, SNPs with missing rate less than 75% before imputation were selected. Divergence times within teosinte and the effective population size of each teosinte were estimated using BPP (A00 model) and MSMC2 (v2.1.1; **Malaspinas AS et al. 2016**). The topological tree in BPP (A00 model) was fixed as the species tree to be the tree with highest posterior probability from the A11 model. Sequences used in the A11 model were applied to estimate the effective population size and divergence time using priors as above. In MSMC2, 2 and 4 haplotype models were applied depending on the number of samples available for each taxon (**Supplementary Table 1**). The mutation rate used in BPP (A00 model) and MSMC2 was 3E-08 (**Clark RM et al. 2005**).

### ABBA-BABA test

We used Patterson’s D statistic (**Green RE et al. 2010; Durand EY et al. 2011**) to test for introgression between teosinte. Assuming *Tripsacum dactyloides* as the outgroup (O), we assessed D statistics for the tree (((H1, H2), H3), O), H1/H2/H3 representing different taxa in *Zea*. The number of ABBA and BABA in each block were calculated in ANGSD (-blockSize 10000). To overcome the problem of nonindependence within the sequence, a block-jackknifing procedure was used to test for statistical significance.

### Linkage disequilibrium, nucleotide diversity and *F*_ST_ calculation

Linkage disequilibrium (*r^2^*) of *mexicana* (81), *parviglumis* (70), *nicaraguensis* (12) and maize (507) were estimated for all bi-allelelic SNPs within 500Kb (-minMAF 0.05 -hwcutoff 0 -missingCutoff 0.5) using Haploview (v4.2; **Barrett JC et al. 2005**). Nucleotide diversity of *mexicana* (81), *parviglumis* (70), *nicaraguensis* (12) and maize (randomly selected 110 individuals) was calculated using ANGSD (v0.930, -doMaf 1 -doMajorMinor 1 -uniqueOnly 1 -minMapQ 30 -minQ 20 -GL 2 -fold 1 -win 5000 -step 5000). Differentiation (*F*_ST_) between maize and teosinte (*mexicana*, *parviglumis*, *nicaraguensis*) was estimated in VCFTools (--fst-window-size 5000; **Danecek P et al. 2011**).

### K-mer analysis

K-mers of teosinte and maize were counted using Jellyfish (v2.3.0; **Marçais G and Kingsford CA 2011**; -m 31). The genome size of *perennis* was set to 5Gb and other species to 2.5Gb. Unique k-mers in each population were obtained with sourmash (v3.2.0; **Brown CT and Irber L. 2016**; --scaled 1000).

### Inversion calling

Genotypes of *mexicana* and *parviglumis* were extracted from the imputed teosinte variation dataset using VCFTools. SNPs with minor allele frequency lower than 0.05 were filtered out and a SNP set was randomly selected to ensure 3 SNPs per kilobase. Inversions were identified using inveRsion (**Cáceres A et al. 2012**) and genotyped by using invClust (**Cáceres A et al. 2015**) with B73 as the reference state. Adjacent inversions were merged to larger regions when the percentage of samples with an inversion was higher than 73% and the distance from each other smaller than 10Mb. The frequency of inversions along chromosomes was plotted using RIdeogram (**Hao Z et al. 2020**). Inversions near the centromeres were also filtered out. GO enrichment analysis of genes located in each inversion region was conducted using PANTHER with default parameters.

### d_N_/d_S_ in teosinte

We randomly selected one individual from each of *parviglumis*, *mexicana*, *huehuetenangensis*, *diploperennis*, *perennis*, *luxurians* and *nicaraguensis* and created a consensus sequence using ANGSD. The branch model in the PAML codeml module (v4.9j; **Yang Z. 2007**) was used to calculate likelihoods of the data under the null hypothesis (H_0_) in which all branches have the same d_N_/d_S_ and the alternative hypothesis (H_a_) in which the test/foreground branch have a d_N_/d_S_ ratio independent from all other/background branches (**Supplementary Figure 13a**). The test tree was fixed as ((((*parviglumis*, *mexicana*), *huehuetenangensis*), (*diploperennis*, *perennis*)), (*luxurians*, *nicaraguensis*)). A likelihood ratio test was used to evaluate the significance of H_a_ over H_0_ (*P*-value < 0.05; **Yang Z. 2007**). Genes related to meiosis were obtained by combining the functionally validated genes in *Arabidopsis thaliana* (**Yant L et al. 2013**). Two large regrowth related QTLs were obtained from Ma et al. (**Ma A et al. 2019**), and genes located in these regions were regarded as candidates related to regrowth. Waterlogging related QTLs were also identified from Guo et al. (**Gou ZF et al. 2021**). GO enrichment analysis of genes with d_N_/d_S_ > 1 (positive selection) and d_N_/d_S_ < 1 (purifying selection) in different tests were conducted using PANTHER with default parameters (**Ashburner M et al. 2000; The Gene Ontology Consortium. 2019**).

### Genome-wide association analysis

Population structure of *mexicana* and *parviglumis* was calculated with admixture (v1.3.0; --cv=10; K=1, 2, 3, 4, 5). The K value with the lowest CV (K=2) was used in downstream analysis. Estimation of the kinship matrix and association analyses using the compressed MLM were performed using TASSEL3 (**Bradbury PJ et al. 2007**), with a *P*-value cut off set to 1/N (N represents the number of SNPs). Latitude, longitude and elevation information were provided by passport data of teosinte accessions from CIMMYT. Phenotypes of global soil properties were extracted from from the Global Soil Dataset for Earth System Modeling (**Wei SG et al. 2014**), a comprehensive Global Soil Dataset for use in Earth System Models by eight layers to the depth of 2.3m (0-0.045, 0.045-0.091, 0.091-0.166, 0.166-0.289, 0.289-0.493, 0.493-0.829, 0.829-1.383 and 1.383-2.296 m) based on the latitude and longitude information of each individual using the R package ncdf4 (http://cirrus.ucsd.edu/~pierce/ncdf/). GWAS was performed on a subset of 29 features identified by hierarchical cluster analysis (**Supplementary Figure 16**).

### Selective sweeps in teosinte and maize

Whole genome scanning for regions of teosinte elevation adaptation and maize temperate adaptation was implemented by a mixed method. First, two genetic maps were obtained from a B73 x Teosinte population (**Liu Z et al. 2016**) and a maize B73 x By804 population (**Pan Q et al. 2016**), and the physical locations were converted to coordinates of the B73 reference sequence, v4 by using CrossMap (**Zhao H et al. 2014**). Second, the genetic distance between SNPs in *mexicana* and *parviglumis* were calculated based on the B73 x Teosinte genetic map, while the distance in temperate maize and tropical maize were calculated based on the B73 x By804 genetic map. Genetic distances between SNPs located between the genetic markers were averaged based on their physical distance. Third, a method based on modeling the likelihood of multi-locus allele frequency differentiation between two tested populations was applied using XP-CLR (**Chen H et al. 2010**; -w1 0.005 100 1000 -p0 0.7) in both the teosinte group (*mexicana*, with *parviglumis* as the reference) and the maize group (temperate maize, with tropical maize as the reference). Finally, a spline-window method GenWin (**Beissinger TM et al. 2015**; smoothness = 100) was applied to smooth the results. The top 5% of the W statistic regions in *parviglumis* and *mexicana* were regarded as candidate teosinte altitude adaptation regions and the top 5% of the W statistic regions in temperate and tropical maize were regarded as candidate maize temperate adaptation regions. Enrichment analysis between inversions and candidate teosinte altitude adaptation regions were conducted by using the shuffle function in BEDTools (v2.25.0; **Quinlan AR and Hall IM. 2010**).

### RNA-seq sampling, library construction and data analysis

The base tissues of V5 stage shoots (1-2 cm) of maize (five tropical maize; five temperate maize) and teosinte (three *parviglumis*; three *mexicana*) were sampled for mRNA and total RNA extraction. Both mRNA and total RNA samples were used for library preparation according to Illumina strand-specific library construction protocols. Libraries were sequenced for both ends using a mixture of platforms (Hi-Seq3000, x10, NovaSeq) with 150 cycles. Considering the coding method for base quality were different in each platform, raw reads were filtered to remove the poor-quality base calls and adaptors specifically for each platform (**Chen S et al. 2018**; NovaSeq: fastp -g -w 2 -l 36; x10: fastp -w 10 -l 36; Hi-Seq3000: Trimmomatic LEADING:3 TRAILING:3 SLIDINGWINDOW:4:15 MINLEN:36). Reads were then aligned to the B73 reference genome (V4) using TopHat2 (v2.2.1; **Kim D et al. 2013**) and read counts for each gene were calculated using htseq-count (**Anders S et al. 2015**). Finally, differentially expressed genes were identified between tropical and temperate maize, as well as between *parviglumis* and *mexicana*, using DESeq2 (**Love MI et al. 2014**) with absolute fold change higher than 1 and *P*-value < 0.05.

### Functional validation of *ZmWOX11*, *ZmCOL9* and *ZmPRR7*

Mutants of *ZmCOL9* and *ZmPRR7* were generated from a high-throughput genome-editing design (**Liu HJ et al. 2020**). In brief, line-specific sgRNAs were filtered based on the assembled pseudo-genome of the receptor KN5585, and a double sgRNAs pool (DSP) approach was used to construct vectors. The vectors were transformed into the receptor KN5585, and the targets of each T_0_ individual were assigned by barcode-based sequencing. The genotype of gene-editing lines was identified by PCR amplification and Sanger sequencing using target-specific primers (**Supplementary Table 23**).

Transgenic lines using DNA fragments of *ZmWOX11* and *ZmCOL9* driven by the *ZmUbi* promoter were created using the modified binary vector pCAMBIA3300. Immature zygotic embryos of maize inbred line KN5585 and hybrid HiII (B73 x A188) were infected with *A. tumefaciens* strain EHA105 harboring the binary vector based on the published method for *ZmWOX11* and *ZmCOL9*, respectively (**Frame BR et al. 2002**). Transgenic plants were identified by qRT-PCR as well as tests for herbicide resistance and the presence of the bar gene.

For stress treatments, maize seeds of the *ZmWOX11* overexpression line were germinated under humid paper in a growth chamber for 7 d at 28□ with 24-h light. The seedlings were then transferred to containers filled with double-distilled water and planted without nutrients for 3 d, followed by nutrient solution for 12 d (normal Hoagland/phosphorus-free Hoagland for Pi stress treatments). In the last 6 d, KH_2_PO_4_ (0, 500μM, 1000μM), PH (control, 4, 8; adjust by using HCl and NaOH), NaCl (0, 100mM, 200mM) and CaCl_2_ (0, 20mM, 40mM) solutions of different concentrations were added to the containers. The solutions were changed every 3 d. Root length of 22-day-old seedlings were measured. Agronomic traits of *ZmWOX11* were investigated in Ezhou province (E114°, N30°) and Hainan province (E109°, N18°). Flowering-time phenotypes of mutants and transgenic plants of *ZmCOL9* and *ZmPRR7* were investigated in Jilin province (E125°, N44°) and Hainan province (E109°, N18°).

## ACKNOWLEDGMENTS

We would like to thank Dr. Suketoshi Taba from CIMMYT for providing teosinte materials. We would like to thank Dr. Jiafa Chen from Henan Agricultural University for providing *Tripsacum dactyloides*. We would like to thank Mr. Hao Liu from the high-throughput computing platform of National Key Laboratory of Crop Genetic Improvement. We would like to thank Andi Kur from North Carolina State University for drawing the picture of teosinte morphological characteristics. This research was supported by the National Key Research and Development Program of China (2020YFE0202300) and National Natural Science Foundation of China (U1901201, 31771879). We would like to acknowledge funding to support this research from the US National Science Foundation grant 1546719 and USDA Hatch project CADPLS2066H to JR-I.

## AUTHOR CONTRIBUTIONS

J.Y., J.R.-I. and N.Y. designed and supervised this study. Y.P., W.L., J.Y., Q.Z., S.G., S.W., Y.W., Y.L., C.J., M.D., M.J., L.J., Y.Y., M.Z. and X.Y. prepared the materials. X.Z. provided the variant calling pipeline. L.C. and J.L. analyzed the data. M.J., X.L., L.C., Y.P., L.Q., Y.Y., X.Y, J.L. and M.Z performed genetic transformation and mutant validation. L.C., M.J., M.H., J.R.-I., N.Y., A.R.F., M.L.W. and J.Y. prepared the manuscript.

## Reference

1. Ray, D. K., Mueller, N. D., West, P. C. & Foley, J. A. Yield trends are insufficient to double global crop production by 2050. PLoS One 8, e66428 (2013).

2. Bailey-Serres, J., Parker, J. E., Ainsworth, E. A., Oldroyd, G. E. D. & Schroeder, J. I. Genetic strategies for improving crop yields. Nature 575, 109–118 (2019).

3. Lesk, C., Rowhani, P. & Ramankutty, N. Influence of extreme weather disasters on global crop production. Nature 529, 84 (2016).

4. Challinor, A. J. et al. A meta-analysis of crop yield under climate change and adaptation. Nature Climate Change 4, 287–291 (2014).

5. Tigchelaar, M., Battisti, D. S., Naylor, R. L. & Ray, D. K. Future warming increases probability of globally synchronized maize production shocks. Proc. Natl. Acad. Sci. USA 115, 6644–6649 (2018).

6. Li, Q. & Yan, J. Sustainable agriculture in the era of omics: knowledge-driven crop breeding. Genome Biol. 21, 154 (2020).

7. Matsuoka, Y. et al. A single domestication for maize shown by multilocus microsatellite genotyping. Proc. Natl. Acad. Sci. USA 99, 6080–6084 (2012).

8. Piperno, D. R., Ranere, A. J., Holst, I., Iriarte. J. & Dickau, R. Starch grain and phytolith evidence for early ninth millennium B.P. maize from the Central Balsas River Valley, Mexico. Proc. Natl. Acad. Sci. USA 106, 5019–5024 (2009).

9. Eyre-Walker, A., Gaut, R. L., Hilton, H., Feldman, D. L. & Gaut, B. S. Investigation of the bottleneck leading to the domestication of maize. Proc. Natl. Acad. Sci. USA 95, 4441–4446 (1998).

10. Tenaillon, M. I., U’Ren, J., Tenaillon, O. & Gaut, B. S. Selection versus demography: a multilocus investigation of the domestication process in maize. Mol. Biol. Evol 21, 1214–1225 (2004).

11. Wright, S. I. et al. The effects of artificial selection on the maize genome. Science 308, 1310–1314 (2005).

12. Beissinger, T. M. et al. Recent demography drives changes in linked selection across the maize genome. Nat. Plants 2, 16084 (2016).

13. Swarts, K. et al. Genomic estimation of complex traits reveals ancient maize adaptation to temperate North America. Science 357, 512–515 (2017).

14. Zampieri, M. et al. When will current climate extremes affecting maize production become the norm? Earth’s Future 7, 113–122 (2019).

15. Zhang, H., Li, Y. & Zhu, J. K. Developing naturally stress-resistant crops for a sustainable agriculture. Nat. Plants. 4, 989–996 (2018).

16. Hufford, M. B., Bilinski, P., Pyhäjärvi, T. & Ross-Ibarra, J. Teosinte as a model system for population and ecological genomics. Trends Genet. 28, 606–615 (2012a).

17. Sánchez González, J. J. et al. Ecogeography of teosinte. PLoS One 13, e0192676 (2018).

18. Mammadov, J. et al. Wild relatives of maize, rice, cotton, and soybean: treasure troves for tolerance to biotic and abiotic Stresses. Front. Plant Sci. 9, 886 (2018).

19. Tian, J. et al. Teosinte ligule allele narrows plant architecture and enhances high-density maize yields. Science 365, 658–664 (2019).

20. Karn, A., Gillman, J. D. & Flint-Garcia, S. A. Genetic analysis of teosinte alleles for kernel composition traits in maize. G3 (Bethesda) 7, 1157–1164 (2017).

21. Li, K. et al. Large-scale metabolite quantitative trait locus analysis provides new insights for high-quality maize improvement. Plant J. 99, 216–230 (2019).

22. Mano, Y., Omori, F. & Takeda, K. Construction of intraspecific linkage maps, detection of a chromosome inversion, and mapping of QTL for constitutive root aerenchyma formation in the teosinte *Zea nicaraguensis*. Mol. Breed 29, 137–146 (2012).

23. Mano, Y. & Omori, F. Flooding tolerance in interspecific introgression lines containing chromosome segments from teosinte (*Zea nicaraguensis*) in maize (*Zea mays* subsp*. mays*). Ann. Bot. 112, 1125–1139 (2013).

24. Lange, E. S. D., Balmer, D., Brigitte, Mauch-Mani. & Turlings, T. C. J. Insect and pathogen attack and resistance in maize and its wild ancestors, the teosintes. New Phytol 204 (2015).

25. Lennon, J. R., Krakowsky, M., Goodman, M., Flint-Garcia, S. & Balint-Kurti, P. J. Identification of alleles conferring resistance to gray leaf spot in maize derived from its wild progenitor species teosinte. Crop Science 56, 222–225 (2015).

26. Lennon, J. R., Krakowsky, M., Goodman, M., Flint-Garcia, S. & Balint-Kurti, P. J. Identification of teosinte alleles for resistance to southern leaf blight in near isogenic maize lines. Crop Science 57, 1973–1983 (2017).

27. Hufford, M. B. et al. The genomic signature of crop-wild introgression in maize. PLoS Genet. 9, e1003477 (2013).

28. Gaut, B. S. & Clegg, M. T. Molecular evolution of the Adh1 locus in the genus *Zea*. Proc. Natl. Acad. Sci. USA 90, 5095–5099 (1993).

29. Hilton, H. & Gaut, B. S. Speciation and domestication in maize and its wild relatives: evidence from the globulin-1 gene. Genetics 150, 863–72 (1998).

30. White, S. E. & Doebley, J. F. The molecular evolution of terminal ear1, a regulatory gene in the genus *Zea*. Genetics 153, 1455–1462 (1999).

31. Ross-Ibarra, J., Tenaillon, M. & Gaut, B. S. Historical divergence and gene flow in the genus *Zea*. Genetics 181, 1399–413 (2009).

32. Wang, Q. & Dooner, H. K. Dynamic evolution of bz orthologous regions in the *Andropogoneae* and other grasses. Plant J. 72, 212–221 (2012).

33. Doebley, J. F. Genetic diversity and population structure of teosinte. Genetics 169, 2241–2254 (2005).

34. Ting Y. C. Spontaneous chromosome inversions of Guatemalan teosintes (*Zea mexicana*). Genetica 36, 229–242 (1965).

35. Ting, Y.C. Common inversion in maize and teosinte. Am. Nat. 101, 87–89 (1967).

36. Ting Y. C. Chromosome polymorphism of teosinte. Genetics 83, 737–742 (1976).

37. Fang, Z. et al. Megabase-scale inversion polymorphism in the wild ancestor of maize. Genetics 191, 883–894 (2012).

38. Pyhäjärvi, T., Hufford, M. B., Mezmouk, S., & Ross-Ibarra, J. Complex patterns of local adaptation in teosinte. Genome Biol. Evol. 5, 1594–609 (2013).

39. Yang, N. et al. Contributions of *Zea mays* subspecies *mexicana* haplotypes to modern maize. Nat. Commun. 8, 1874 (2017).

40. Lamb, J. C. & Birchler, J. A. Retroelement genome painting: cytological visualization of retroelement expansions in the genera *Zea* and *Tripsacum*. Genetics 173, 1007–21 (2006).

41. Tenaillon, M. I., Hufford, M. B., Gaut, B. S. & Ross-Ibarra, J. Genome size and transposable element content as determined by high-throughput sequencing in maize and *Zea luxurians*. Genome Biol. Evol. 3, 219–29 (2011).

42. Chia JM et al. Maize HapMap2 identifies extant variation from a genome in flux. Nat Genet. 44, 803–807 (2012).

43. Fustier, M. A. et al. Common gardens in teosintes reveal the establishment of a syndrome of adaptation to altitude. PLoS Genet. 15, e1008512 (2019).

44. Janzen, G. M. et al. Demonstration of local adaptation of maize landraces by reciprocal transplantation. bioRxiv (2021).

45. Fustier M. A. et al. Signatures of local adaptation in lowland and highland teosintes from whole-genome sequencing of pooled samples. Mol. Ecol. 26, 2738–2756 (2017).

46. Aguirre-Liguori, J. A. et al. Divergence with gene flow is driven by local adaptation to temperature and soil phosphorus concentration in teosinte subspecies (*Zea mays parviglumis* and *Zea mays mexicana*). Mol. Ecol. 28, 2814–2830 (2019).

47. Yang, X. H. et al. Characterization of a global germplasm collection and its potential utilization for analysis of complex quantitative traits in maize. Mol Breeding. 28, 511–526 (2011).

48. Flouri, T., Jiao, X., Rannala, B. & Yang, Z. Species tree inference with BPP using genomic sequences and the multispecies coalescent. Mol. Biol. Evol. 35, 2585–2593 (2018).

49. Lee, T. H., Guo, H., Wang, X., Kim, C. & Paterson, A. H. SNPhylo: a pipeline to construct a phylogenetic tree from huge SNP data. BMC Genomics 15, 162 (2014).

50. Wang, X. et al. Genome alignment spanning major Poaceae lineages reveals heterogeneous evolutionary rates and alters inferred dates for key evolutionary events. Mol Plant. 8, 885–898 (2015).

51. Buckler, E. S. 4th. & Holtsford, T. P. Zea systematics: ribosomal ITS evidence. Mol Biol Evol. 13, 612–622 (1996).

52. Malaspinas, A. S. et al. A genomic history of Aboriginal Australia. Nature 538, 207–214 (2016).

53. Warburton, M. L. et al. Gene flow among different teosinte taxa and into the domesticated maize gene pool. Genet Resour Crop Evol 58, 1243–1261 (2011).

54. Janzen, G. M., Wang, L. & Hufford, M. B. The extent of adaptive wild introgression in crops. New Phytol. 221, 1279–1288 (2019).

55. van Heerwaarden, J., Hufford, M. B. & Ross-Ibarra, J. Historical genomics of North American maize. Proc. Natl. Acad. Sci. USA 109, 12420–12425 (2012).

56. Bellon, M. R. et al. Evolutionary and food supply implications of ongoing maize domestication by Mexican *campesinos*. Proc Biol Sci. 285, 20181049 (2018).

57. Faria, R., Johannesson, K., Butlin, R. K. & Westram, A. M. Evolving Inversions. Trends Ecol Evol. 34, 239–248 (2019).

58. Crow, T. et al. Gene regulatory effects of a large chromosomal inversion in highland maize. PLoS Genet. 16, e1009213 (2020).

59. Cáceres, A. & González, J. R. Following the footprints of polymorphic inversions on SNP data: from detection to association tests. Nucleic Acids Res. 43, e53 (2015)

60. Pawlowski, W. P. et al. Coordination of meiotic recombination, pairing, and synapsis by PHS1. Science 303, 89–92 (2004).

61. Ronceret, A., Doutriaux, M. P., Golubovskaya, I. N. & Pawlowski, W. P. PHS1 regulates meiotic recombination and homologous chromosome pairing by controlling the transport of RAD50 to the nucleus. Proc. Natl. Acad. Sci. USA 106, 20121–20126 (2009).

62. Ting, Y. C. Chromosomes of maize-teosinte hybrids. Cambridge (MA) The Bussey Institution of Harvard University (1964).

63. Jha, P., Ochatt, S. J. & Kumar, V. WUSCHEL: a master regulator in plant growth signaling. Plant Cell Rep 39, 431–444 (2020).

64. Wei S., Dai Y., Duan Q., Liu B. & Hua Y. A global soil data set for earth system modeling. J. Adv. Model. Earth Syst. 6, 249–263 (2014).

65. Chen, Z. H., Nimmo, G. A., Jenkins, G. I. & Nimmo, H. G. BHLH32 modulates several biochemical and morphological processes that respond to Pi starvation in *Arabidopsis*. Biochem J. 405, 191–198 (405).

66. Verma, V., Ravindran, P. & Kumar, P. P. Plant hormone-mediated regulation of stress responses. BMC Plant Biol. 16, 86 (2016).

67. Ku, Y. S., Sintaha, M., Cheung, M. Y. & Lam, H. M. Plant Hormone Signaling Crosstalks between Biotic and Abiotic Stress Responses. Int. J. Mol. Sci. 19, 3206 (2018).

68. Hoopes, G. M. et al. An updated gene atlas for maize reveals organ-specific and stress-induced genes. Plant J. 97, 1154–1167 (2019).

69. Merrill, W. L et al. The diffusion of maize to the southwestern United States and its impact. Proc. Natl. Acad. Sci. USA 106, 21019–21026 (2009).

70. Hart, J. B. & Hetty, L. R. Extending the phytolith evidence for early maize (*Zea mays* ssp. mays) and squash (*Cucurbita* sp.) in Central New York. Am. Antiq. 72, 563–583 (2007).

71. Yan, L et al. Parallels between natural selection in the cold-adapted crop-wild relative *Tripsacum dactyloides* and artificial selection in temperate adapted maize. Plant J. 99, 965–977 (2019).

72. Buckler, E. S. et al. The genetic architecture of maize flowering time. Science 325, 714–718 (2009).

73. Han, J. J., Jackson, D. & Martienssen, R. Pod corn is caused by rearrangement at the tunicate1 locus. Plant Cell 24, 2733–2744 (2012).

74. Liu, H. et al. High-Throughput CRISPR/Cas9 Mutagenesis Streamlines Trait Gene Identification in Maize. Plant Cell 32, 1397–1413 (2020).

75. Hufford, M. B., Martínez-Meyer, E., Gaut, B. S., Eguiarte, L. E. & Tenaillon, M. I. Inferences from the historical distribution of wild and domesticated maize provide ecological and evolutionary insight. PLoS One 7, e47659 (2012b).

76. Aguirre-Liguori, J. A. et al. Connecting genomic patterns of local adaptation and niche suitability in teosintes. Mol. Ecol. 26, 4226–4240 (2017).

77. Gui, S. et al. ZEAMAP, a comprehensive database adapted to the maize multiomics era. iScience 23, 101241 (2020).

78. Kumar, M. Impact of climate change on crop yield and role of model for achieving food security. Environ. Monit. Assess. 188, 465 (2016).

79. Xu, H., Twine, T. E. & Girvetz, E. Climate change and maize yield in Iowa. PLoS One 11, e0156083 (2016).

80. Giannini, T. C. et al. Projected climate change threatens pollinators and crop production in Brazil. PLoS One 12, e0182274 (2017).

81. Meyer, R. S. & Purugganan, M. D. Evolution of crop species: genetics of domestication and diversification. Nat. Rev. Genet. 14, 840–852 (2013).

82. Kirkpatrick, M. & Barton, N. Chromosome inversions, local adaptation and speciation. Genetics 173, 419–34 (2006).

83. Lorant, A. et al. The potential role of genetic assimilation during maize domestication. PLoS One 12, e0184202 (2017).

84. Hufford, M. B. et al. Comparative population genomics of maize domestication and improvement. Nat Genet. 44, 808–11 (2012c).

85. Suzuki, N. Hormone signaling pathways under stress combinations. Plant Signal. Behav. 11, e1247139 (2016).

86. Miller, R. N., Costa Alves, G. S. & Van Sluys, M. A. Plant immunity: unravelling the complexity of plant responses to biotic stresses. Ann. Bot. 119, 681–687 (2017).

87. Yang, J. et al. The crosstalks between jasmonic acid and other plant hormone signaling highlight the involvement of jasmonic acid as a core component in plant response to biotic and abiotic stresses. Front. Plant Sci. 10, 1349 (2019).

88. O’Brien, A. M., Sawers, R. J. H., Strauss, S. Y., & Ross-Ibarra, J. Adaptive phenotypic divergence in an annual grass differs across biotic contexts. Evolution 73, 2230–2246 (2019).

89. Fernie, A. R., & Yan, J. Targeting key genes to tailor old and new crops for a greener agriculture. Mol Plant. 13, 354–356 (2020).

## Reference

90. Bolger, A. M., Lohse, M. & Usadel, B. Trimmomatic: a flexible trimmer for Illumina sequence data. Bioinformatics 30, 2114–20 (2014).

91. Jiao Y et al. Improved maize reference genome with single-molecule technologies. Nature 546, 524–527 (2017).

92. Langmead, B. & Salzberg, S. L. Fast gapped-read alignment with Bowtie 2. Nat. Methods. 9, 357–359 (2012).

93. Li, H. et al. The Sequence Alignment/Map format and SAMtools. Bioinformatics 25, 2078–2079 (2009).

94. Chang, C. C. et al. Second-generation PLINK: rising to the challenge of larger and richer datasets. Gigascience 4, 7 (2015).

95. Alexander, D. H., Novembre, J. & Lange, K. Fast model-based estimation of ancestry in unrelated individuals. Genome Res. 19, 1655–1664 (2009).

96. Yang, J., Lee, S. H., Goddard, M. E., & Visscher, P. M. GCTA: a tool for genome-wide complex trait analysis. Am. J. Hum. Genet. 88, 76–82 (2011).

97. Letunic, I. & Bork, P. Interactive tree of life (iTOL) v4: recent updates and new developments. Nucleic Acids Res. 47, W256–W259 (2019).

98. Korneliussen, T. S., Albrechtsen, A. & Nielsen, R. ANGSD: Analysis of Next Generation Sequencing Data. BMC Bioinform. 15, 356 (2014).

99. Bouckaert, R. R. DensiTree: making sense of sets of phylogenetic trees. Bioinformatics 26, 1372–1373 (2010).

100. Browning, S. R. & Browning, B. L. Rapid and accurate haplotype phasing and missing-data inference for whole-genome association studies by use of localized haplotype clustering. Am. J. Hum. Genet. 81, 1084–1097 (2007).

101. Clark, R. M., Tavaré, S. & Doebley, J. Estimating a nucleotide substitution rate for maize from polymorphism at a major domestication locus. Mol Biol Evol. 22, 2304–2312 (2005).

102. Green, R. E. et al. A draft sequence of the Neandertal genome. Science 328, 710–722 (2010).

103. Durand, E. Y., Patterson, N., Reich, D. & Slatkin, M. Testing for ancient admixture between closely related populations. Mol Biol Evol. 28, 2239–2252 (2011).

104. Barrett, J. C., Fry, B., Maller, J. & Daly, M. J. Haploview: analysis and visualization of LD and haplotype maps. Bioinformatics 21, 263–265 (2005).

105. Danecek, P. et al. The variant call format and VCFtools. Bioinformatics 27, 2156–2158 (2011).

106. Marçais, G. & Kingsford, C. A fast, lock-free approach for efficient parallel counting of occurrences of k-mers. Bioinformatics 27, 764–770 (2011).

107. Brown, C. T. & Irber, L. sourmash: a library for MinHash sketching of DNA. J. Open Source Softw. 1, 27 (2016).

108. Cáceres, A., Sindi, S. S., Raphael, B. J., Cáceres, M. & González, J. R. Identification of polymorphic inversions from genotypes. BMC Bioinformatics. 13, 28 (2012).

109. Hao, Z. et al. RIdeogram: drawing SVG graphics to visualize and map genome-wide data on the idiograms. PeerJ Comput. Sci. 6, e251 (2020).

110. Yang, Z. PAML 4: phylogenetic analysis by maximum likelihood. Mol. Biol. Evol. 24, 1586–1591 (2007).

111. Yant, L. et al. Meiotic adaptation to genome duplication in *Arabidopsis arenosa*. Curr Biol. 23, 2151–2156 (2013).

112. Ma, A. et al. The genetics and genome-wide screening of regrowth loci, a key component of perennialism in *Zea diploperennis*. G3 (Bethesda) 9, 1393–1403 (2019).

113. Guo, Z. et al. Identification of major QTL for waterlogging tolerance in maize using genome-wide association study and bulked sample analysis. J Appl Genet. Epub ahead of print. (2021).

114. Ashburner, M. et al. Gene ontology: tool for the unification of biology. The Gene Ontology Consortium. Nat. Genet. 25, 25–29 (2000).

115. The Gene Ontology Consortium. The Gene Ontology Resource: 20 years and still GOing strong. Nucleic Acids Res. 47, D330–D338 (2019).

116. Bradbury, P. J. et al. TASSEL: software for association mapping of complex traits in diverse samples. Bioinformatics 23, 2633–2635 (2007).

117. Liu, Z. et al. Expanding Maize Genetic Resources with Predomestication Alleles: Maize-Teosinte Introgression Populations. Plant Genome 9 (2016).

118. Pan, Q. et al. Genome-wide recombination dynamics are associated with phenotypic variation in maize. New Phytol. 210, 1083–1094 (2016).

119. Zhao, H. et al. CrossMap: a versatile tool for coordinate conversion between genome assemblies. Bioinformatics. 30, 1006–1007 (2014).

120. Chen, H., Patterson, N. & Reich, D. Population differentiation as a test for selective sweeps. Genome Res. 20, 393–402 (2010).

121. Beissinger, T. M., Rosa, G. J., Kaeppler, S. M., Gianola, D. & de Leon, N. Defining window-boundaries for genomic analyses using smoothing spline techniques. Genet. Sel. Evol. 47, 30 (2015).

122. Quinlan, A. R. & Hall, I. M. BEDTools: a flexible suite of utilities for comparing genomic features. Bioinformatics. 26, 841–842 (2010).

123. Chen, S., Zhou, Y., Chen, Y. & Gu, J. fastp: an ultra-fast all-in-one FASTQ preprocessor. Bioinformatics 34, i884–i890 (2018).

124. Kim, D. et al. TopHat2: accurate alignment of transcriptomes in the presence of insertions, deletions and gene fusions. Genome Biol. 14, R36 (2013).

125. Anders, S., Pyl, P. T. & Huber, W. HTSeq--a Python framework to work with high-throughput sequencing data. Bioinformatics 31, 166–169 (2015).

126. Love, M. I., Huber, W. & Anders, S. Moderated estimation of fold change and dispersion for RNA-seq data with DESeq2. Genome Biol. 15, 550 (2014).

127. Frame, B. R. et al. Agrobacterium tumefaciens-mediated transformation of maize embryos using a standard binary vector system. Plant Physiol. 129, 13–22 (2002).

