## Supplementary Figures and Tables for "Portrait of a genus: the genetic diversity of *Zea*"

**Supplementary Figure 1.** The minor allele frequency (MAF) distribution of *Zea* haplotype map.

**Supplementary Figure 2.** STRUCTURE and principle component analysis (PCA) of 507 maize and 183 teosinte.

**Supplementary Figure 3.** Phylogenetic tree of *Zea* genus.

**Supplementary Figure 4.** A11 species tree from the 10,000,000 Markov Chain Monte Carlo (MCMC).

**Supplementary Figure 5.** Population split time inference.

**Supplementary Figure 6.** ABBA-BABA analysis of introgression.

**Supplementary Figure 7.** The distribution of average nucleotide diversity.

**Supplementary Figure 8.** Private SNPs and InDels in *Zea*.

**Supplementary Figure 9.** LD, $F_{ST}$ and nucleotide diversity of *nicaraguensis*, *mexicana*, *parviglumis* and maize.

**Supplementary Figure 10.** Frequency of inversions in *parviglumis* and *mexicana*.

**Supplementary Figure 11.** Principal component analysis and haplotype of 14 large inversions.

**Supplementary Figure 12.** GO and pathway enrichment analysis in inversions.

**Supplementary Figure 13.** Genes that under positive and purifying selection in teosinte.

**Supplementary Figure 14.** GWAS identification of a candidate gene for highland adaptation.

**Supplementary Figure 15.** The response of *ZmWOX11* with different iron treatments.

**Supplementary Figure 16.** GWAS identification of candidate gene for soil properties.

**Supplementary Figure 17.** Manhattan plot of the representative soil properties.

**Supplementary Figure 18.** Enrichment analysis between 14 large inversions and candidate altitude adaptation windows from XP-CLR.

**Supplementary Figure 19.** Plant hormone signal transduction pathway from KEGG.

**Supplementary Figure 20.** Comparison between genes under selection and without selection.

**Supplementary Figure 21.** Phenotype analysis of different CRISPR/Cas9 mutation *ZmCOL9* in different environments.

**Supplementary Figure 22.** Phenotype analysis of different overexpression lines for *ZmCOL9* in different environments.

**Supplementary Figure 23.** Phenotype analysis of CRISPR/Cas9 mutation for *ZmPRR7* in different environments.

**Supplementary Tables**

**Supplementary Table 8.** Large inversions in *parviglumis* and *mexicana*.

**Supplementary Table 23.** Primers used in transgenic study.

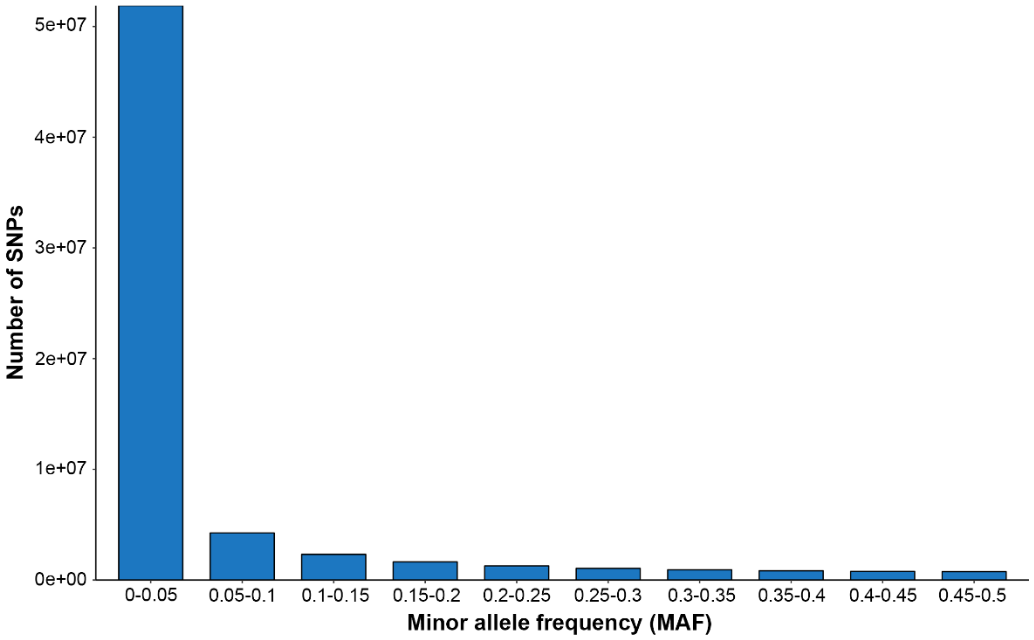


**Supplementary Figure 1. The minor allele frequency (MAF) distribution of *Zea* haplotype map.** Each bar in the plot indicates the number of SNPs in each MAF bin.

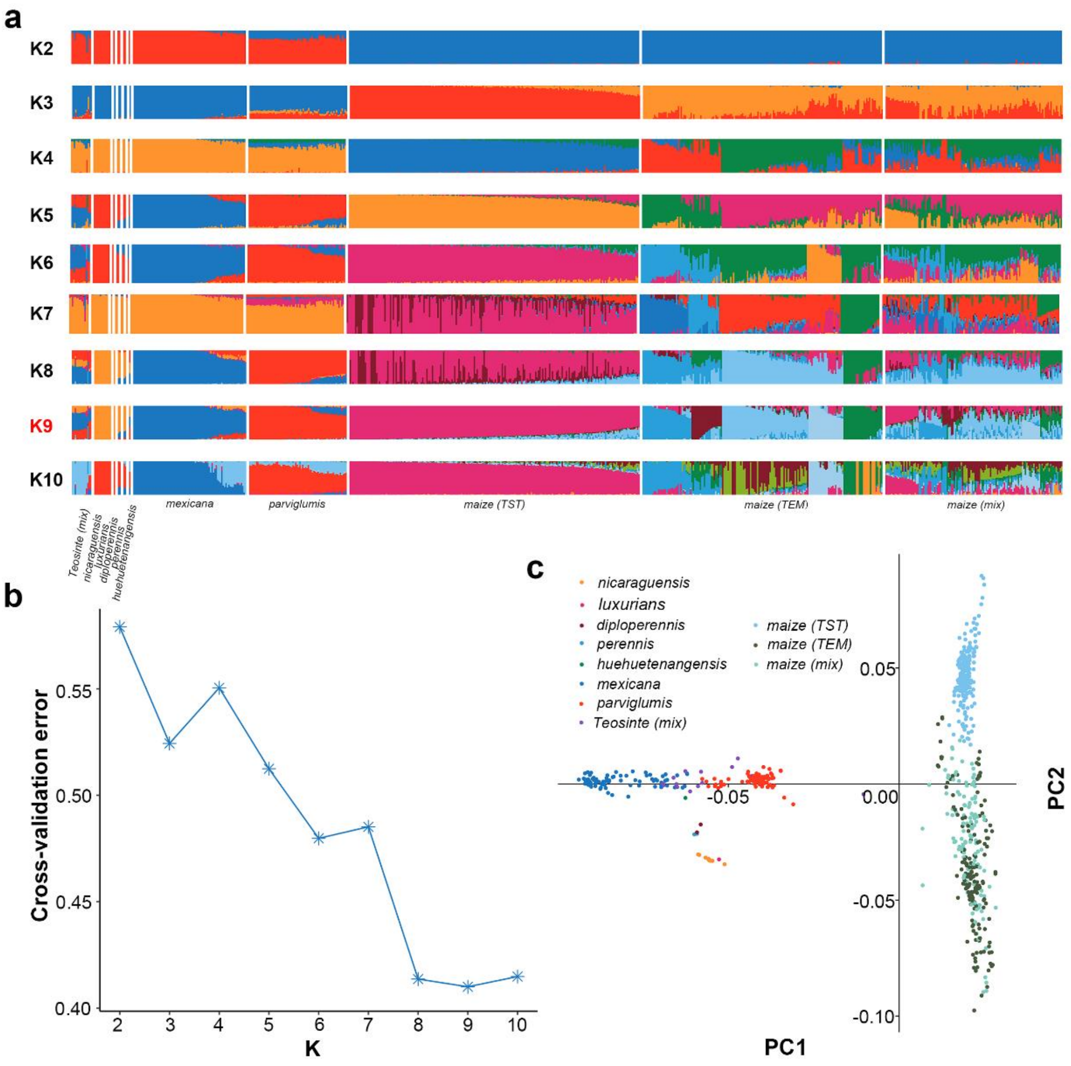


**Supplementary Figure 2. STRUCTURE and principle component analysis (PCA) of 507 maize and 183 teosinte. a,** Population structure for K=2-10. maize (TEM) indicates temperate maize, maize (TST) indicates tropical maize. **b**, Cross-validation error in K=2-10, K=9 with lowest cross validation error. **c**, PCA of maize and teosinte; points are colored according to the admixture result (K=9).

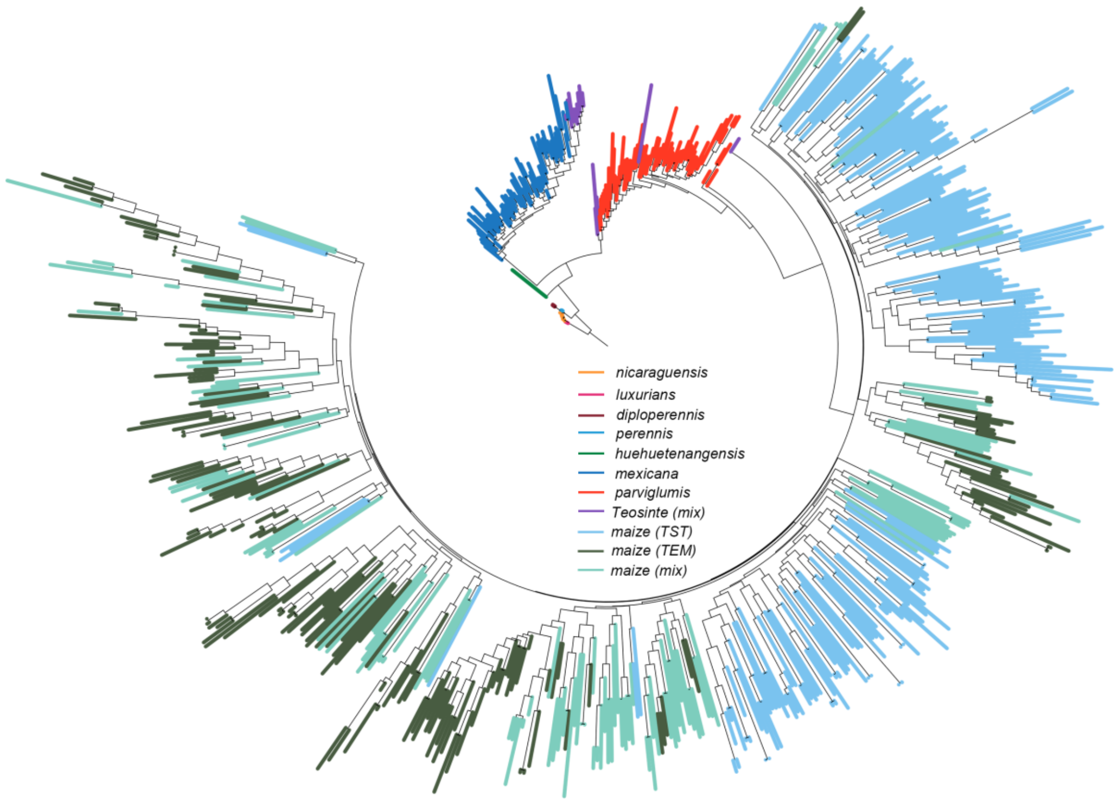


**Supplementary Figure 3. Phylogenetic tree of *Zea* genus**. ML tree was conducted with SNPhylo (Lee TH et al. 2014). Populations are colored based on the admixture result for K=9.

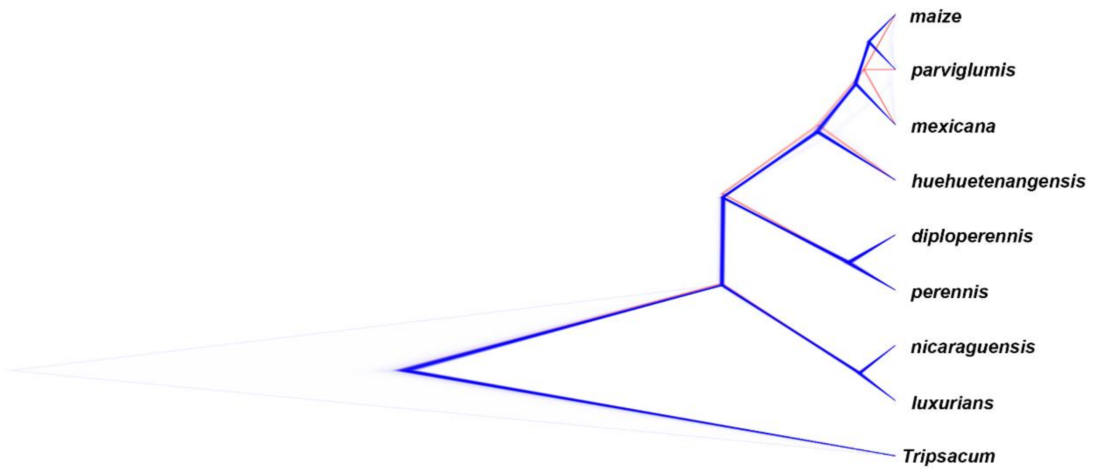


**Supplementary Figure 4. A11 species tree from the 10,000,000 Markov chain Monte Carlo (MCMC) simulation.** DensiTree (Bouckaert RR. 2010) plotted only showing the consensus trees (downsampling to 1,000,000 MCMC generations).

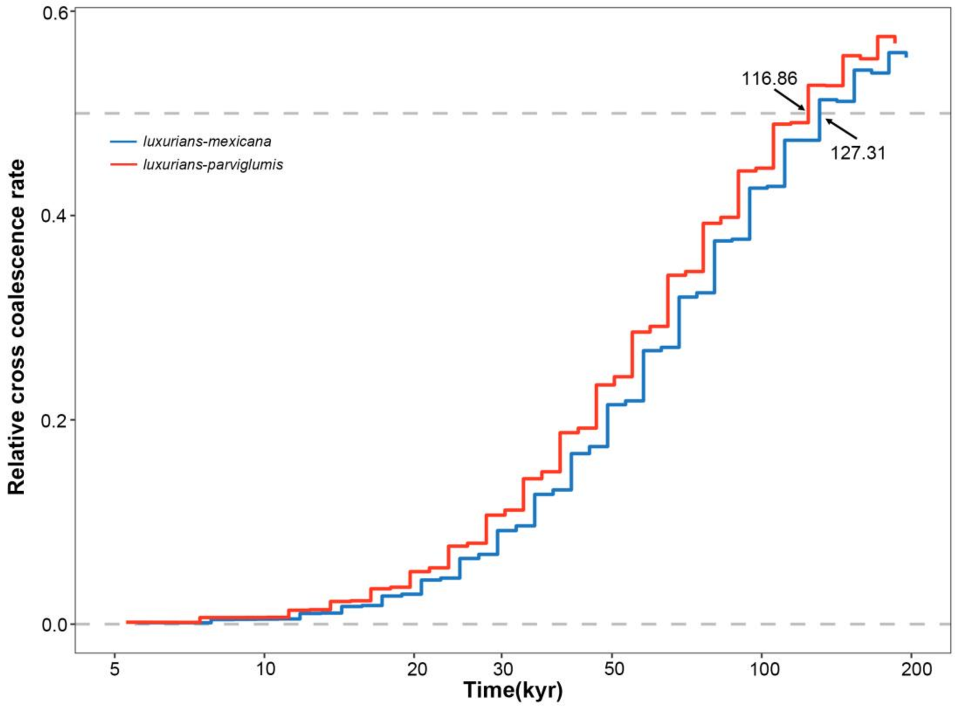


**Supplementary Figure 5. Population split time inference.** MSMC2 (Malaspinas AS et al. 2016) cross-coalescence results for *luxurians* (2 haplotypes) compared to *mexicana* and *parviglumis* (4 haplotypes).

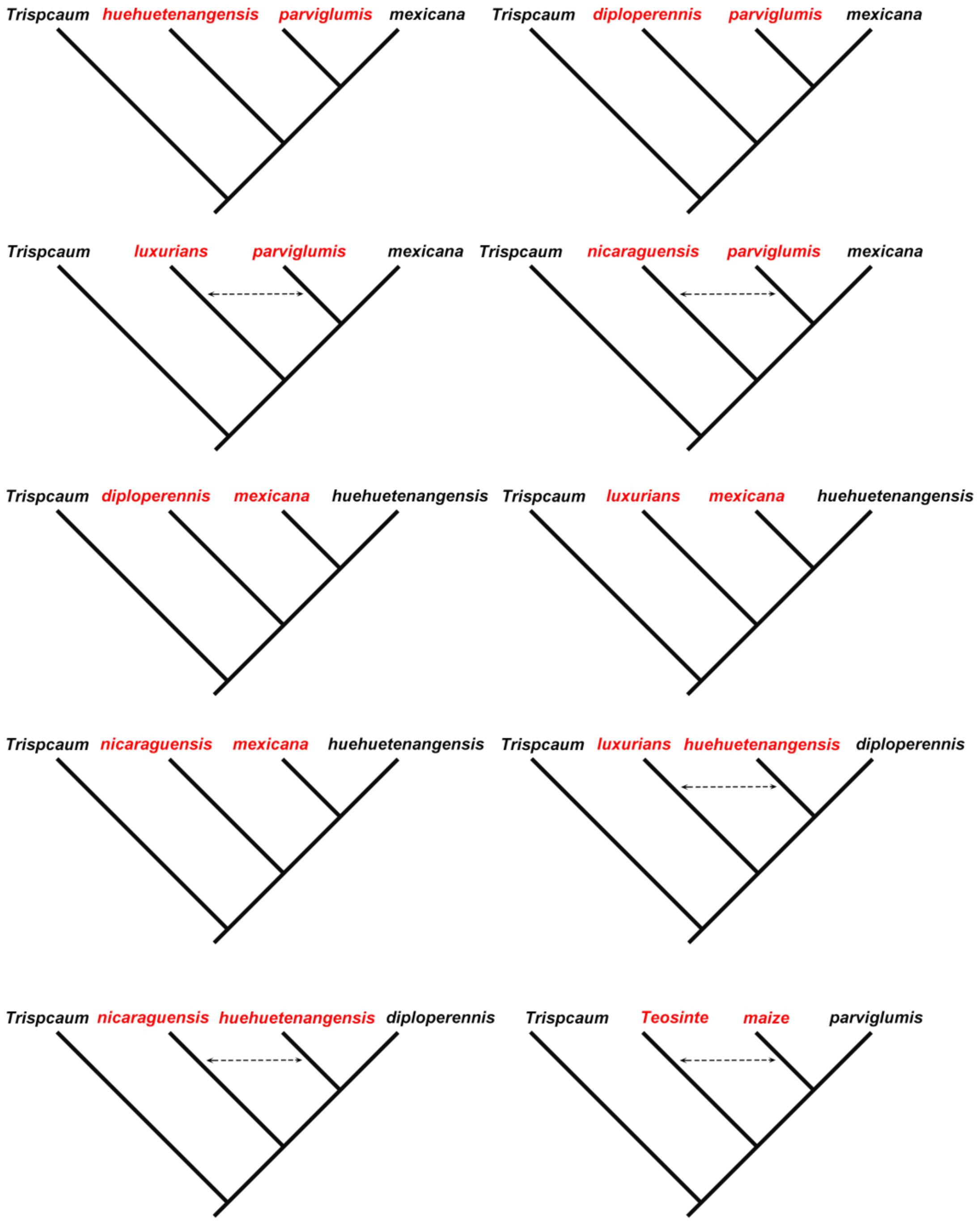


**Supplementary Figure 6. ABBA-BABA analysis of introgression.** Four-species trees used in ABBA-BABA analyses among *Zea* genus. The dashed arrow indicates a positive D statistic. “Teosinte” in the last subplot indicates separate tests for *mexicana* and *huehuetenangensis*.

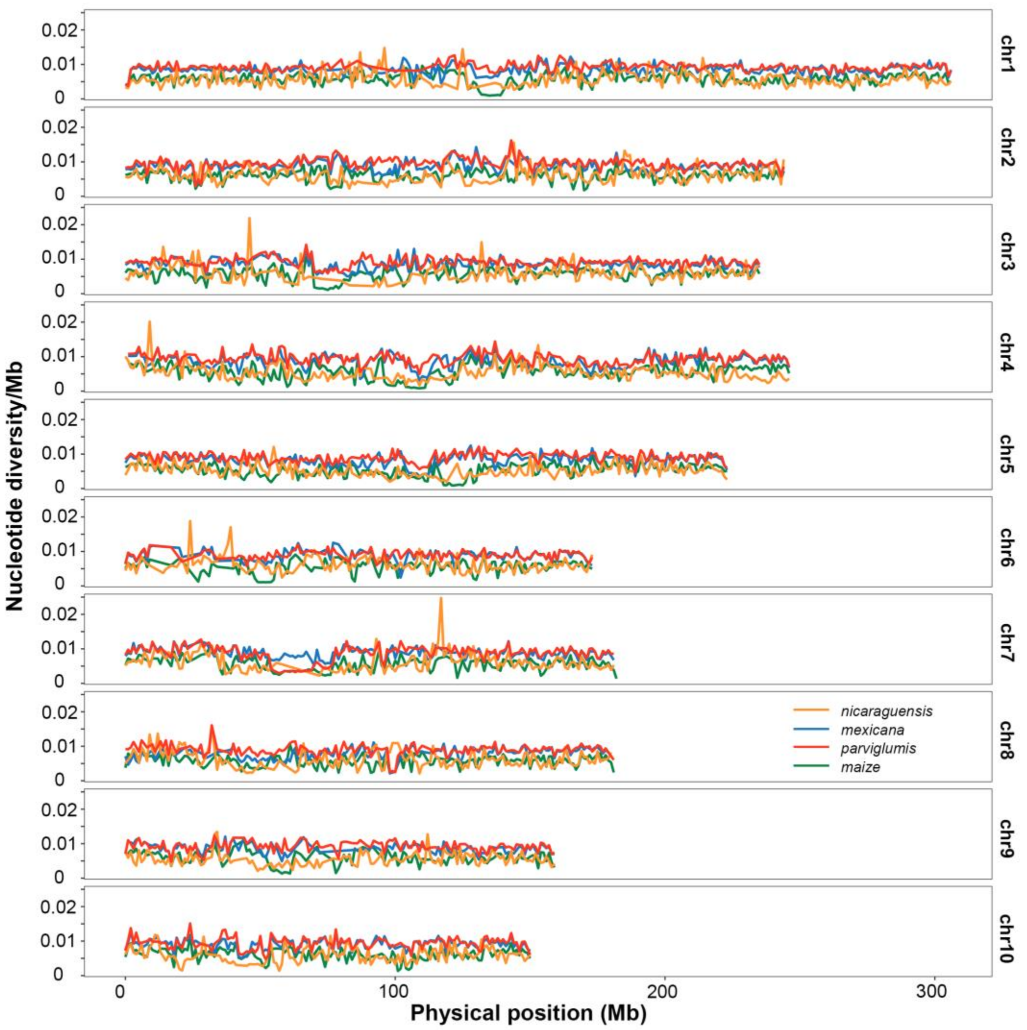


**Supplementary Figure 7. The distribution of average nucleotide diversity.** Line plot of four species that have sufficient sample size (n>=12) for nucleotide diversity analysis. Cultivated maize was down-sampled to 110 randomly selected individuals.

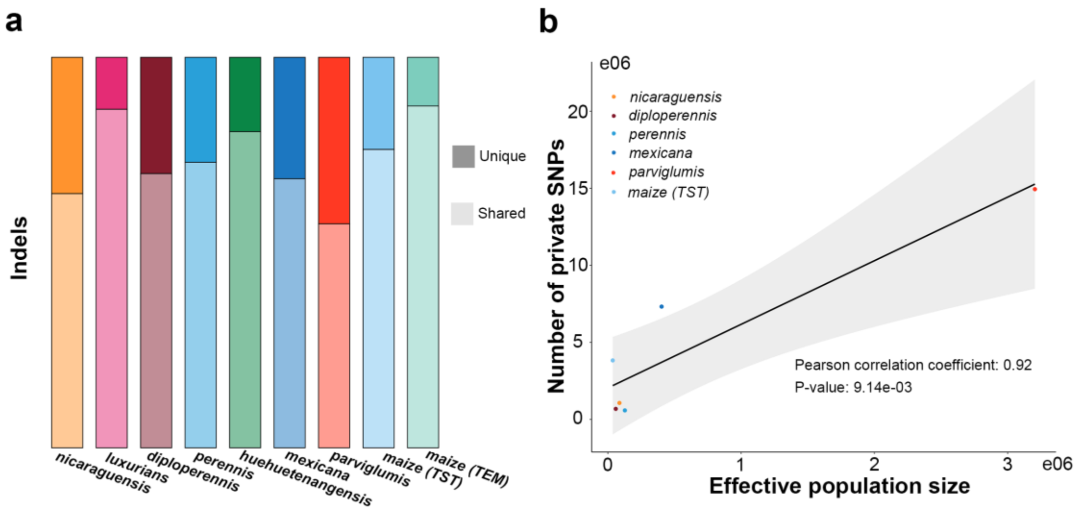


**Supplementary Figure 8. Private SNPs and InDels in *Zea*. a**, The private InDels in *Zea*. **b**, Correlation between effective population size and private SNPs.

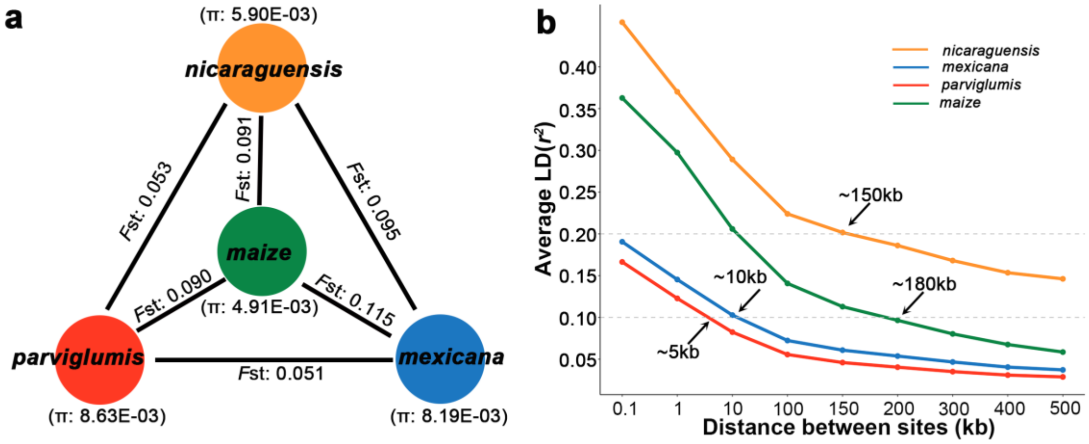


**Supplementary Figure 9. LD, $F_{ST}$ and nucleotide diversity of *nicaraguensis*, *mexicana*, *parviglumis* and maize. a,** Mean nucleotide diversity in each population and mean population differentiation $F_{ST}$ between populations. **b,** LD decay of *nicaraguensis*, *mexicana*, *parviglumis*, *mexicana*. Labels indicate the distance at which mean $r^2 = 0.1$ or $r^2 = 0.2$.

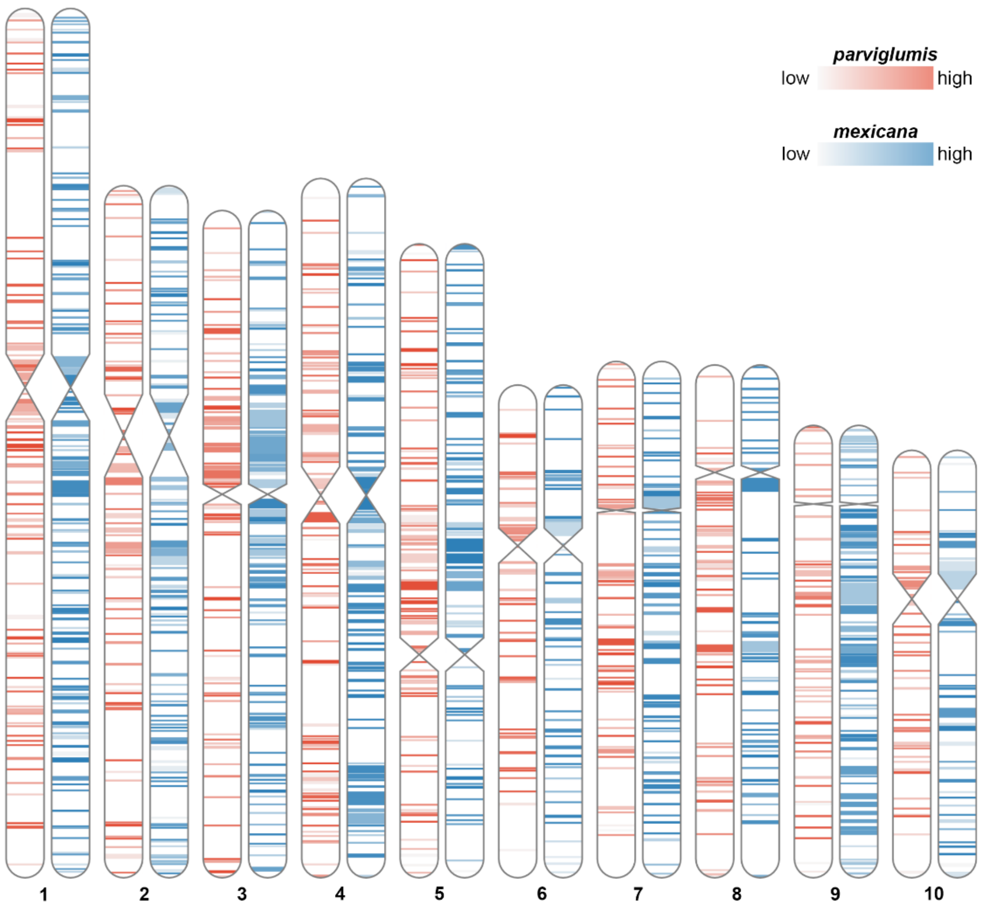


**Supplementary Figure 10. Frequency of inversions in *parviglumis* and *mexicana*.** The karyotype shows for each chromosome a heatmap of the percentage of samples with an inversion.

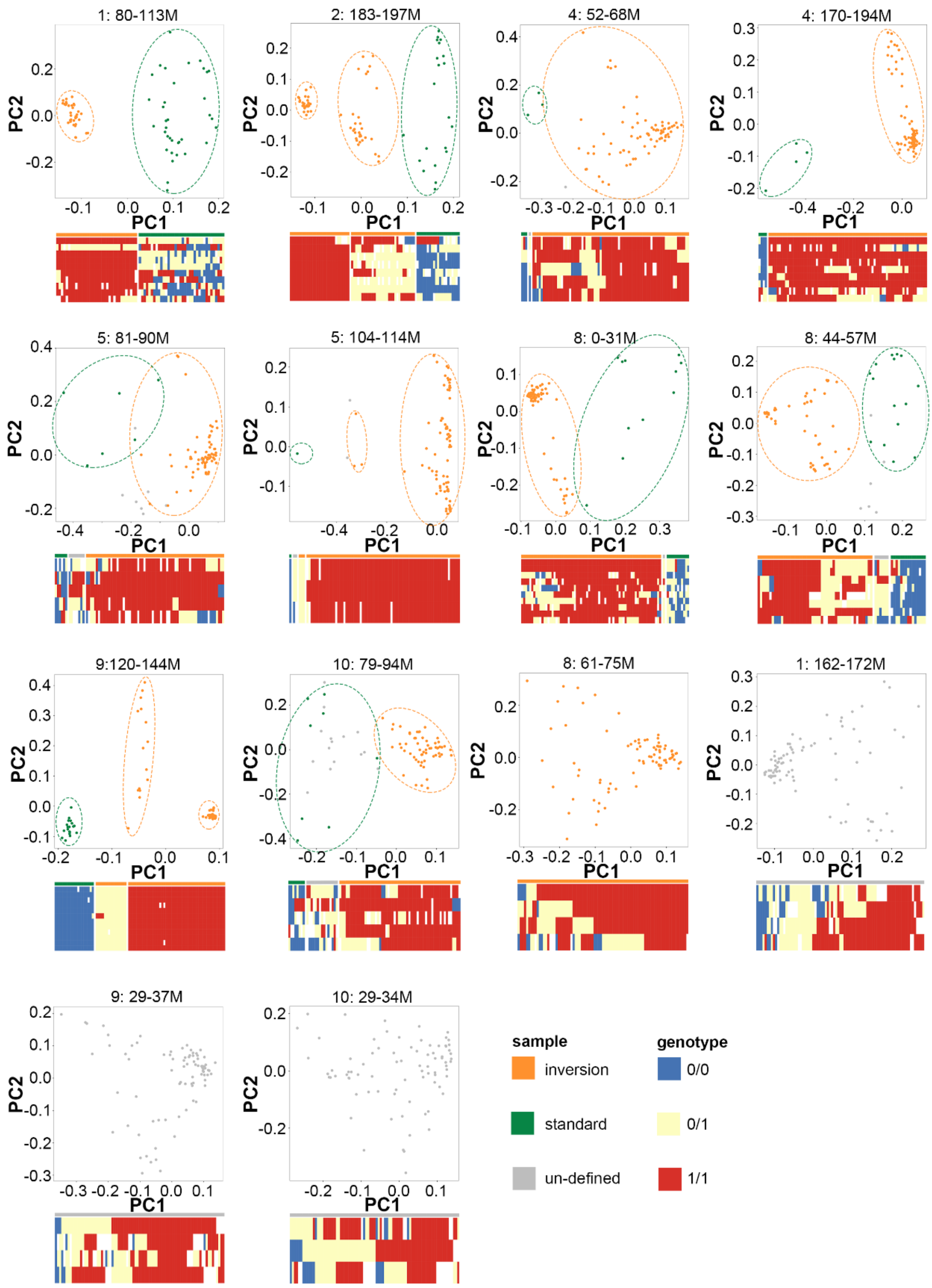


**Supplementary Figure 11. Principal component analysis and haplotype of 14 large inversions.** Dotplot shows the principal component analysis of SNPs in the inversion region. Each point represents a sample. Heatmap shows the genotype of inversion identified by invClust. Bar behind the heatmap shows the sample type.

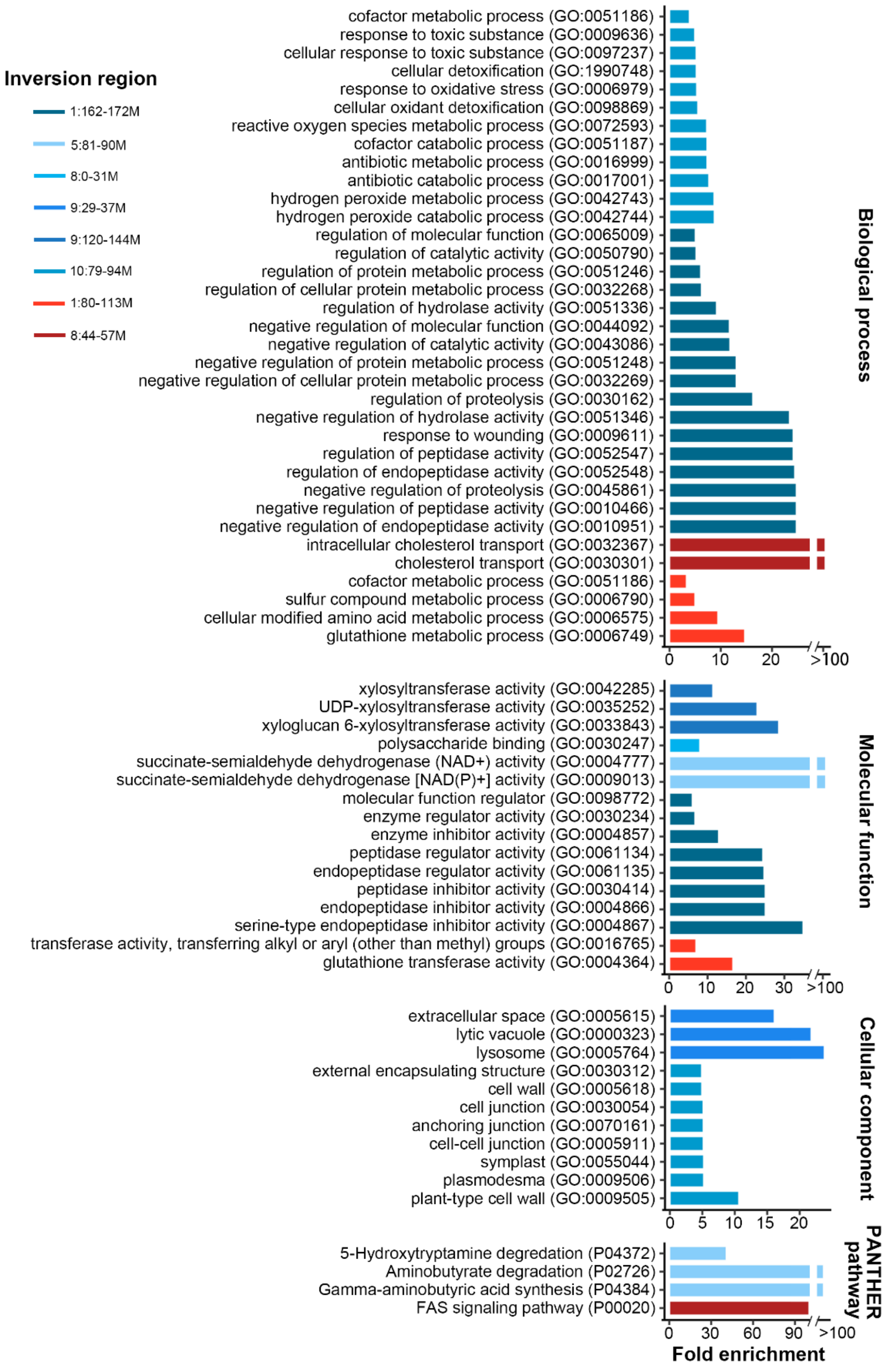


**Supplementary Figure 12. GO and pathway enrichment analysis in inversions.** Bar plot indicated the fold change of genes compared to the background.

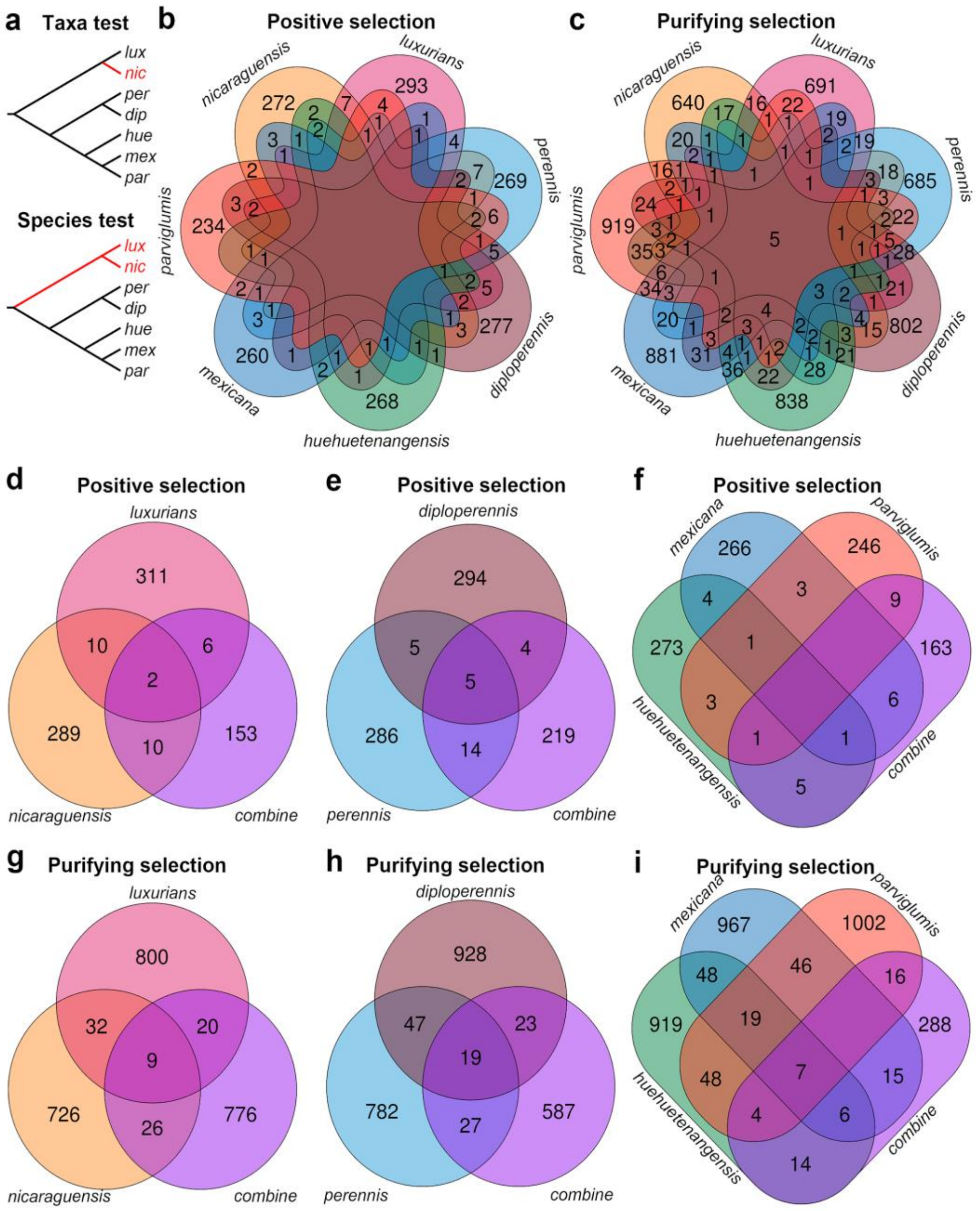


**Supplementary Figure 13. Genes under positive and purifying selection in teosinte.** **a**, Phylogenies used for selection tests, one example for taxa selection, and one example for species selection (test/foreground branch was colored by red). *lux*: *luxurians*, *nic*: *nicaraguensis*, *per*: *perennis*, *dip*: *diploperennis*, *hue*: *huehuetenangensis*, *mex*: *mexicana*, *par*: *parviglumis*. **b-i**, Numbers in the VENN plot indicated genes under positive or purifying selection in that taxa or species (regard as test/foreground branch, other teosinte as background branches). "combine" indicates genes under positive or purifying selection in waterlogging species (*nicaraguensis* and *luxurians*), perennial species (*diploperennis* and *perennis*) and *Zea mays* (*huehuetenangensis*, *mexicana* and *parviglumis*).

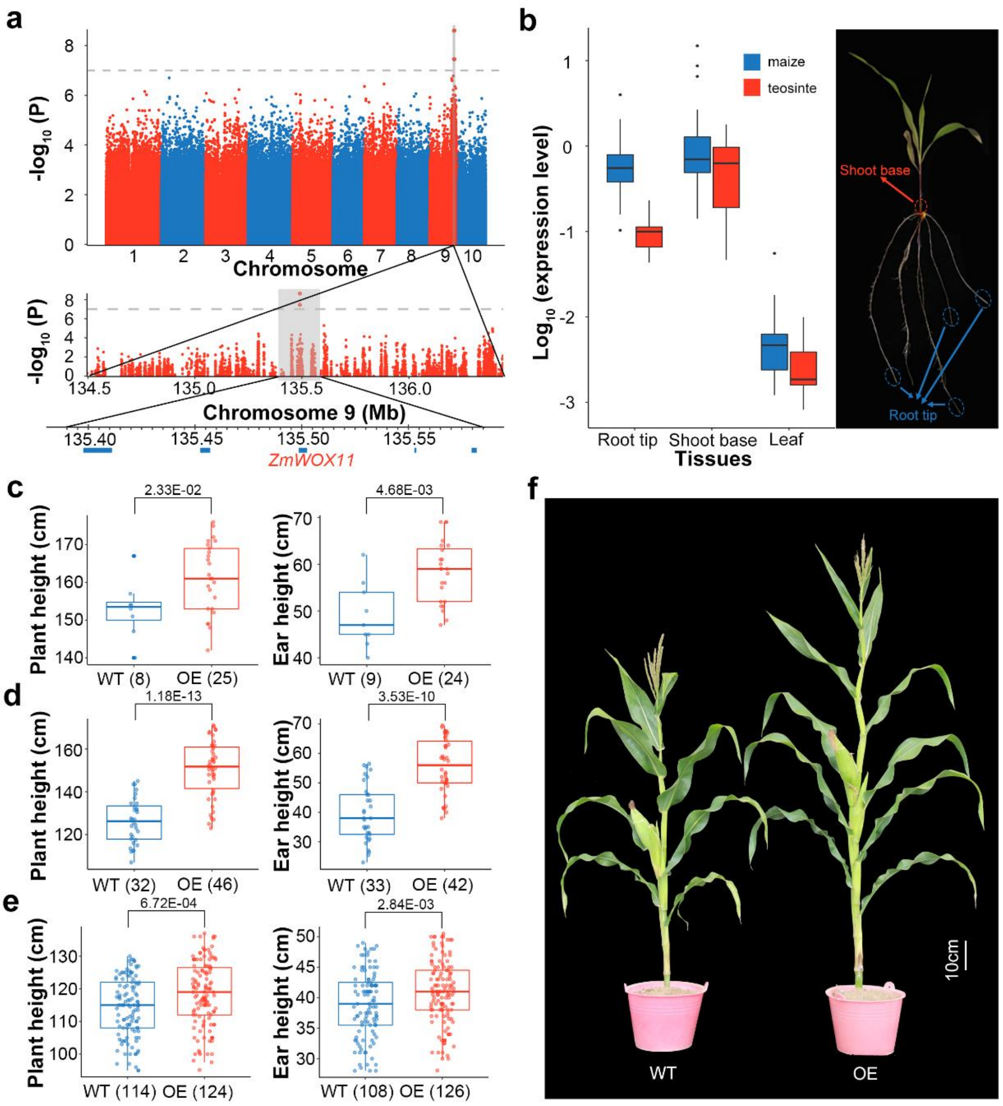


**Supplementary Figure 14. GWAS identification of a candidate gene for highland adaptation. a,** Manhattan plot of the whole genome (top), 134.5Mb-136.5Mb of chromosome 9 (middle), and five genes in the 100kb around the peak SNP (bottom). **b**, RT-qPCR analysis of *ZmWOX11* expression in tips of secondary root (1-2cm), shoot base (1-2cm) and leaf of maize and teosinte. Expression levels are relative to the root of B73 (which regard as 1). The circles in the right picture indicate the tissues harvested for RNA extraction (Supplementary Table 14). **c**, **d, e** Plant and ear height in (**c**) Hainan province (2019; China; E109°, N18°; WT: wild type), (**d**) Hainan province (2020; China; E109°, N18°; WT: wild type) and (**e**) Ezhou province (2020; China; E114°, N30°); OE: overexpression line; ns: no significance). Traits with z-score higher than 1.5 were removed. Two-tailed t-test *P*-value shown. **f**, Image of plant height and ear height of wild type and overexpression line of *ZmWOX11* (2020; Hainan; 89 days after seeding).

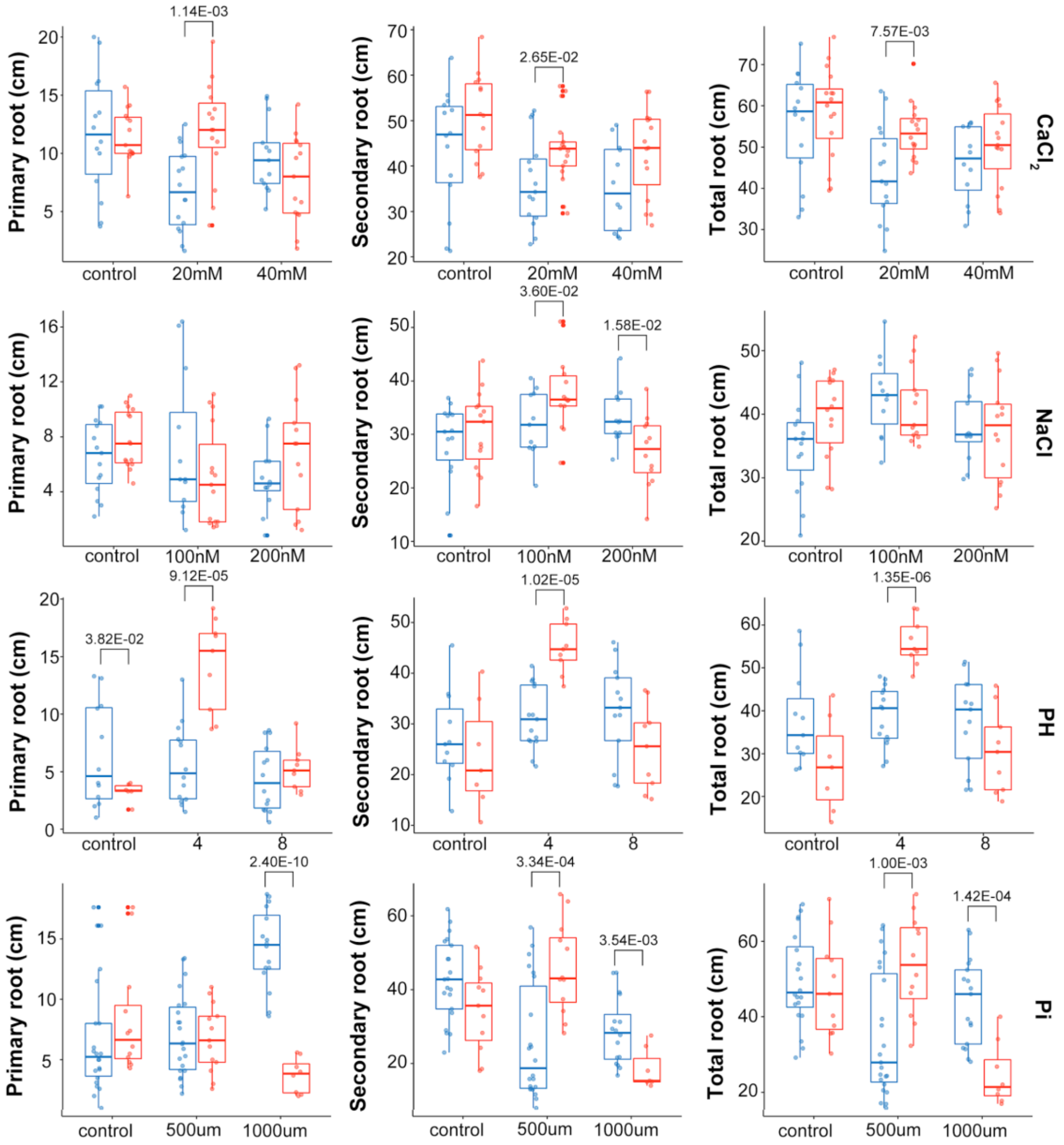


**Supplementary Figure 15. The response of *ZmWOX11* with different iron treatments.** Root length of twenty-two days old KN5585 and *ZmWOX11* overexpression lines (KN5585 background) grown in Hoagland solution (control) and treatment with different iron treatment for 12 days. Traits with z-score higher than 1.5 were removed.

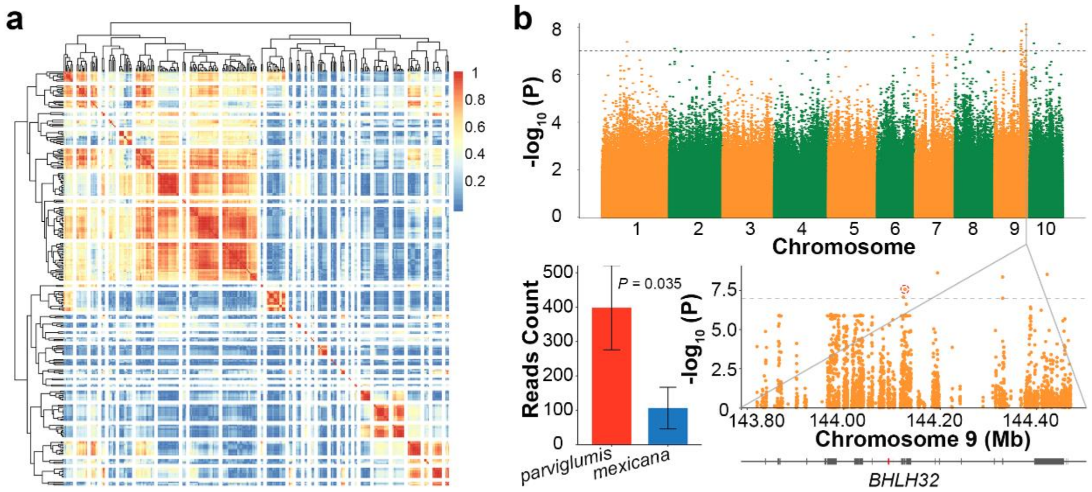


**Supplementary Figure 16. GWAS identification of candidate gene for soil properties**. **a**, Heatmap shows the pearson correlation coefficient between 243 soil properties. The gaps indicate the clusters (n=29). **b**, eGWAS identification of candidate genes. Manhattan plot of the whole genome for electrical conductivity under 0-0.045m (top) and 143.80Mb-144.40Mb of chromosome 9 (right bottom). Point that circled in Manhattan plot show the significant SNP that located closest to the candidate gene (right bottom). RNA-seq read count for *Zm00001d047878* of *parviglumis* and *mexicana* in shoot base (left bottom).

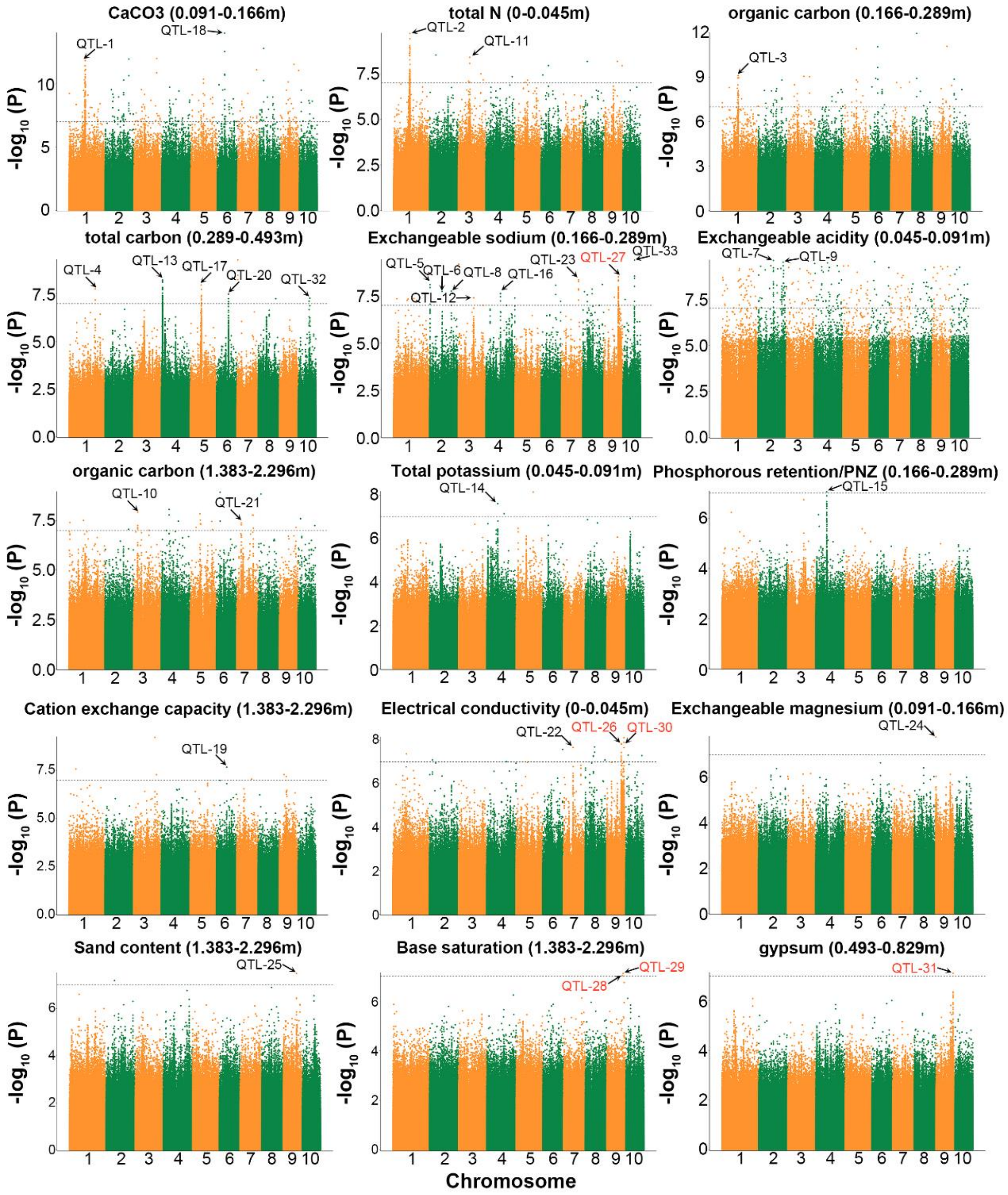


**Supplementary Figure 17. Manhattan plot of the representative soil properties.** Detailed information of QTLs are listed in Supplementary Table 16. QTLs located in *Inv9e* are marked with red color.

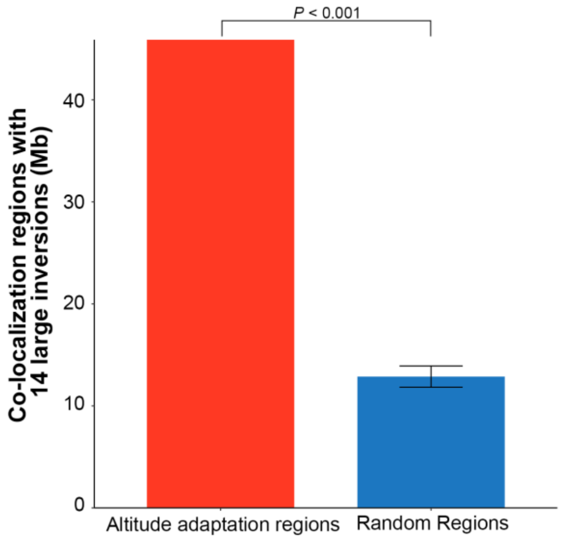


**Supplementary Figure 18. Enrichment analysis between 14 large inversions and candidate altitude adaptation windows from XP-CLR.** *P*-value was calculated under 1,000 simulations.

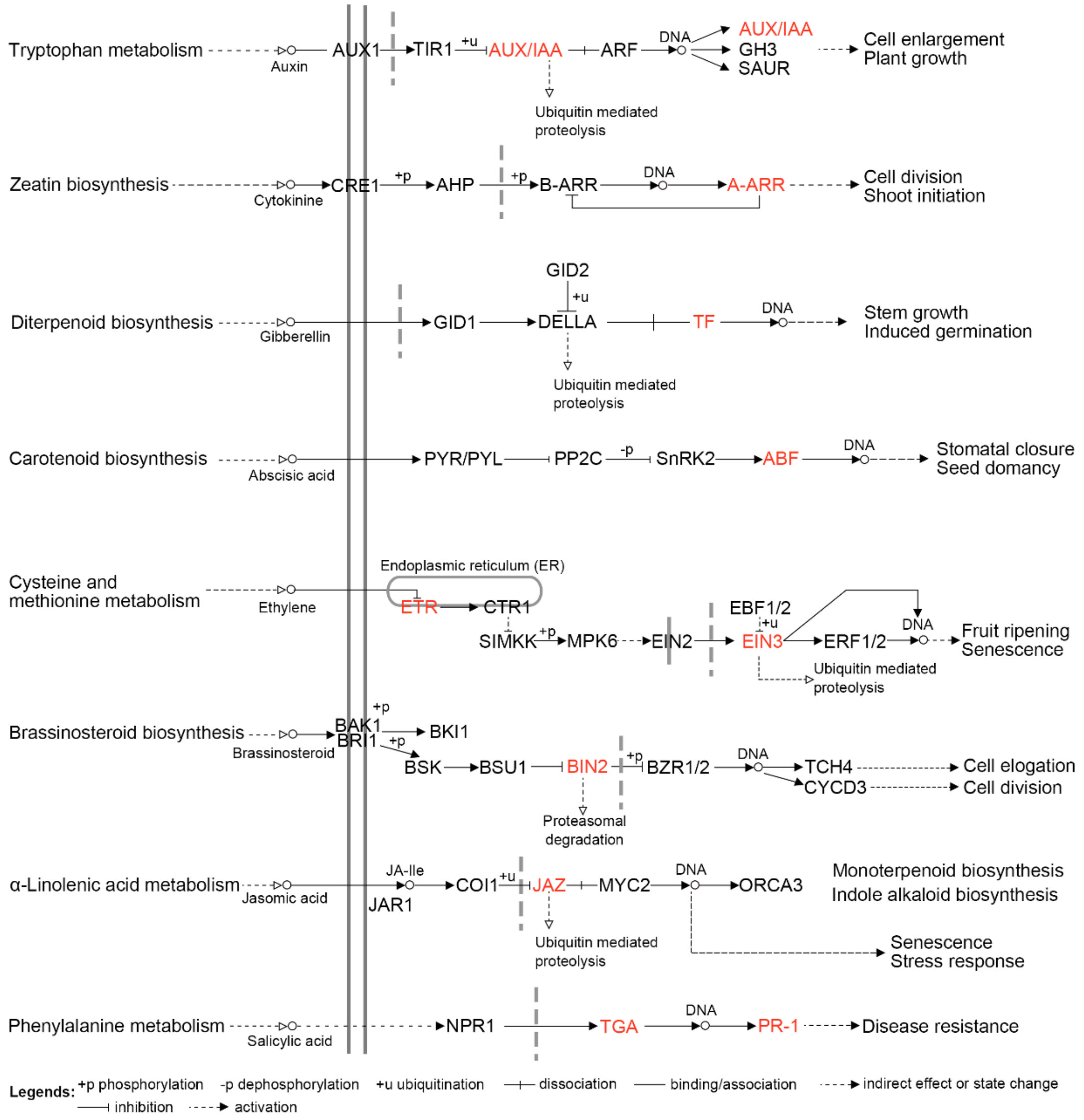


**Supplementary Figure 19. Plant hormone signal transduction pathway from KEGG.** Candidate altitude adaptation genes that were annotated in plant hormone signal transduction pathway by KEGG were colored by red.

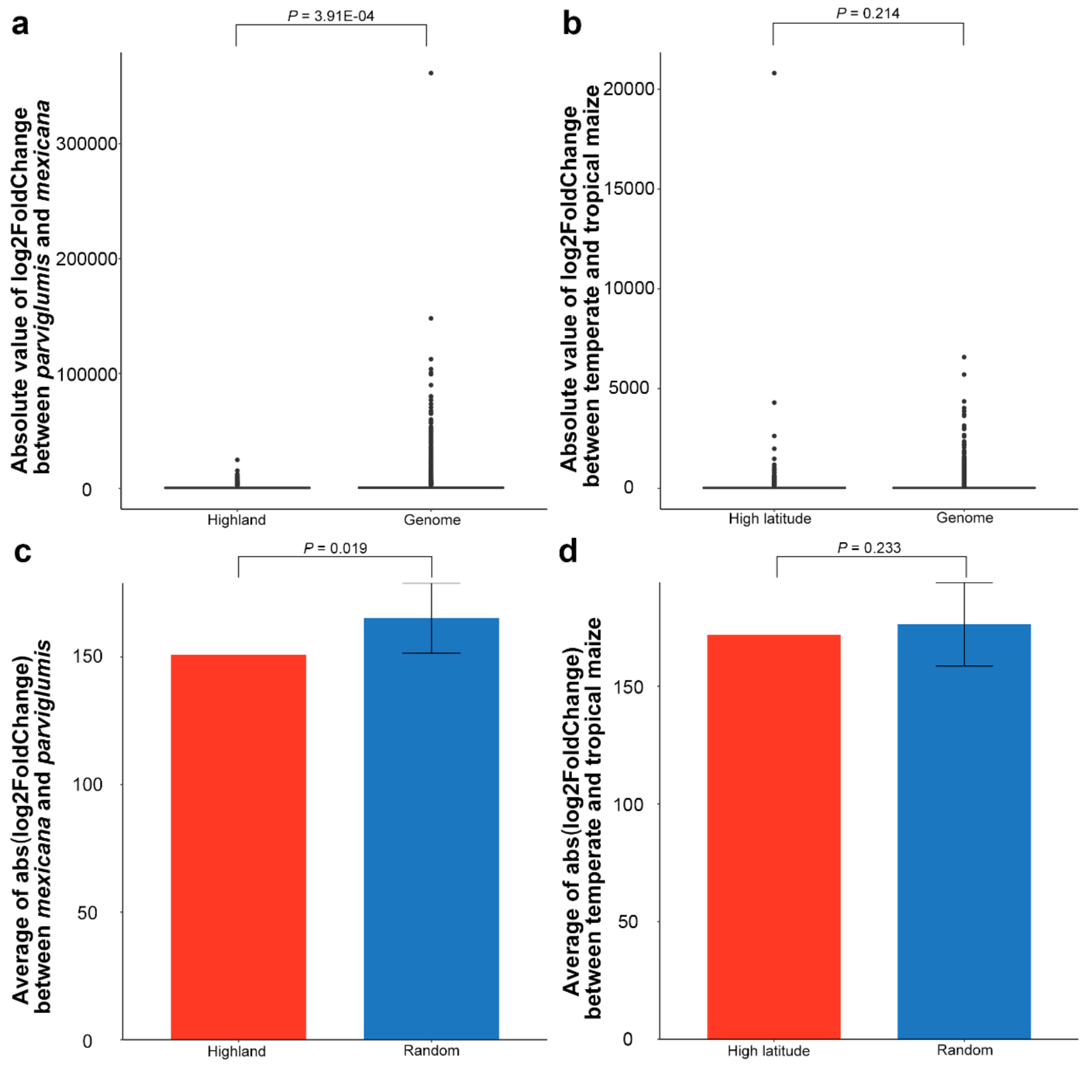


**Supplementary Figure 20**. **Comparison between genes under selection and without selection.** Box plot shows the absolute log2 fold change of gene expressions between **a**, *mexicana* and *parviglumis*. **b**, temperate and tropical maize. *P*-value was calculated by performing Welch Two Sample t-test. Bar plot shows the mean value of absolute log2 fold change of gene expressions between **c**, *mexicana* and *parviglumis* and **d**, temperate and tropical maize. The error bar shows the standard deviation, *P*-value was calculated using 1,000 simulations.

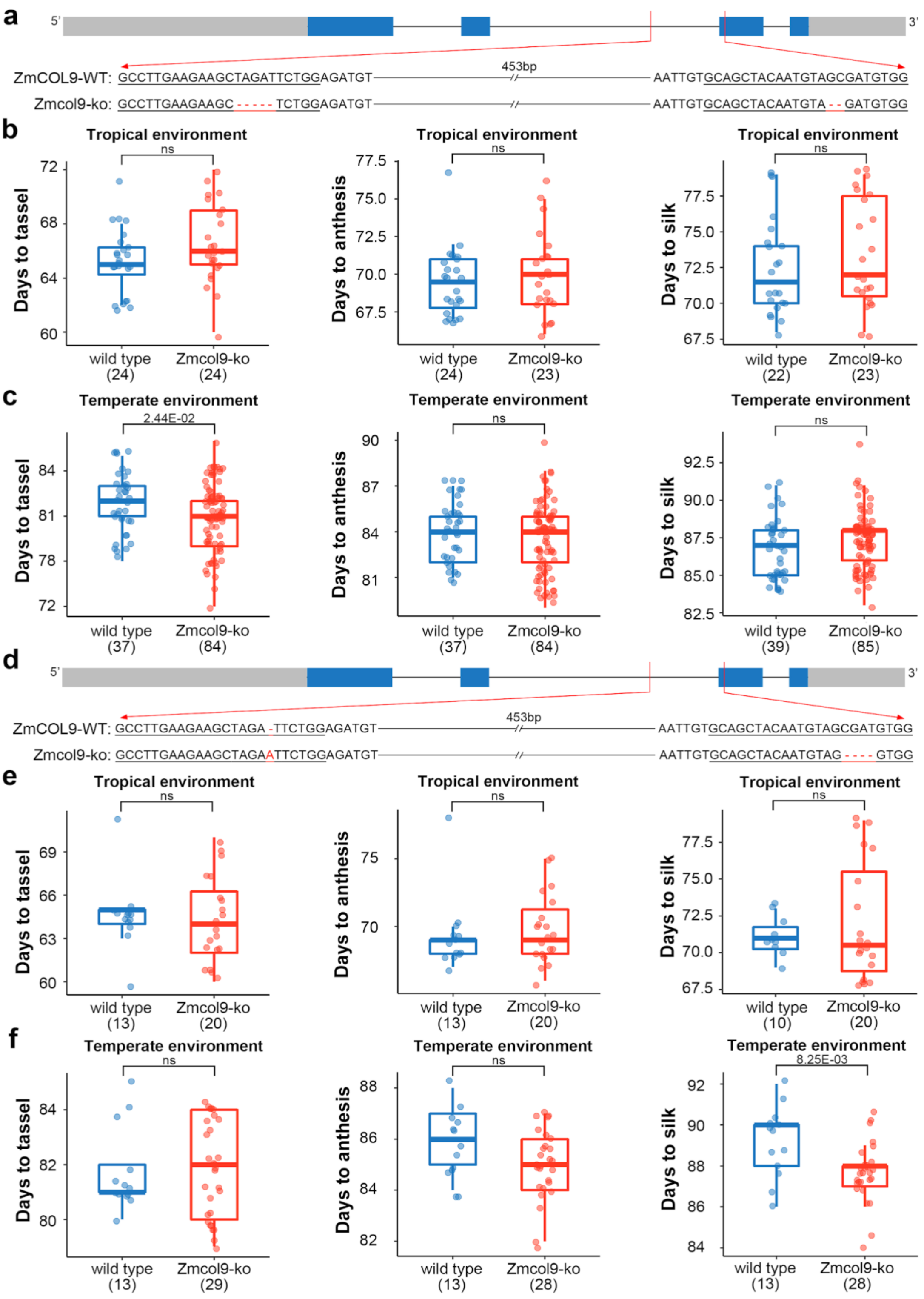


**Supplementary Figure 21. Phenotype analysis of different CRISPR/Cas9 mutation *ZmCOL9* in different environments. a,** Gene structure and sequences of *ZmCOL9* target regions in wild type, Zmcol9 CRISPR/Cas9 knockout mutants 1. **b,** Statics of days to tassel, days to anthesis, days to silk in Hainan province (2019; China; E109°, N18°; CRISPR/Cas9 mutation). **c,** Statics of days to tassel, days to anthesis, days to silk in Jilin province (2020; China; E125°, N44°; CRISPR/Cas9 mutation). **d,** Gene structure and sequences of *ZmCOL9* target regions in wild type, Zmcol9 CRISPR/Cas9 knockout mutants 2. **e,** Statics of days to tassel, days to anthesis, days to silk in Hainan province (2019; China; E109°, N18°; CRISPR/Cas9 mutation). **f,** Statics of days to tassel, days to anthesis, days to silk in Jilin province (2020; China; E125°, N44°; CRISPR/Cas9 mutation).

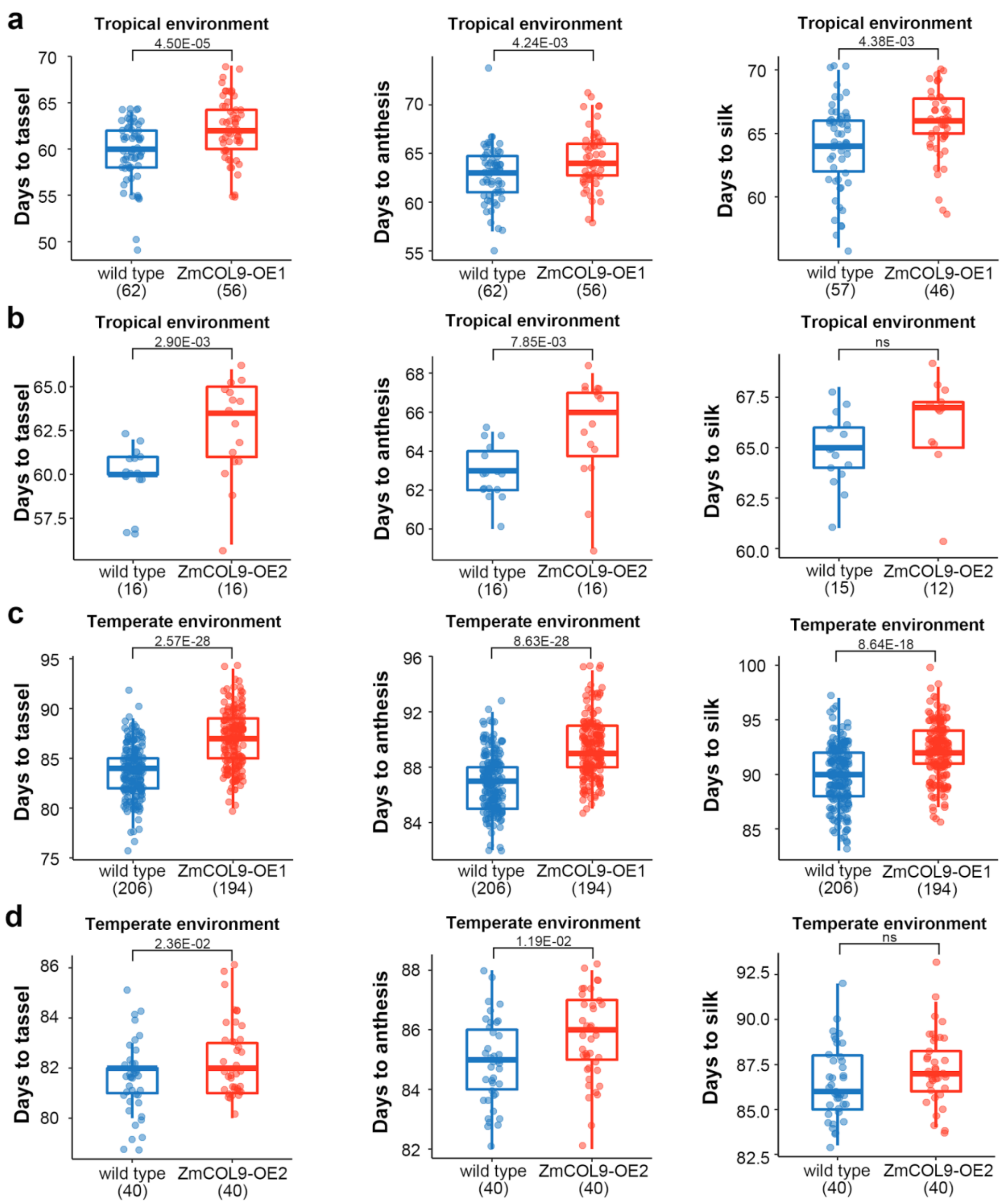


**Supplementary Figure 22. Phenotype analysis of different overexpression lines for *ZmCOL9* in different environments. a,** Statics of days to tassel, days to anthesis, days to silk in Hainan province (2019; China; E109°, N18°; overexpression line 1). **b,** Statics of days to tassel, days to anthesis, days to silk in Hainan province (2019; China; E109°, N18°; overexpression line 2). **c,** Statics of days to tassel, days to anthesis, days to silk in Jilin province (2020; China; E125°, N44°; overexpression line 1). **d,** Statics of days to tassel, days to anthesis, days to silk in Jilin province (2020; China; E125°, N44°; overexpression line 2).

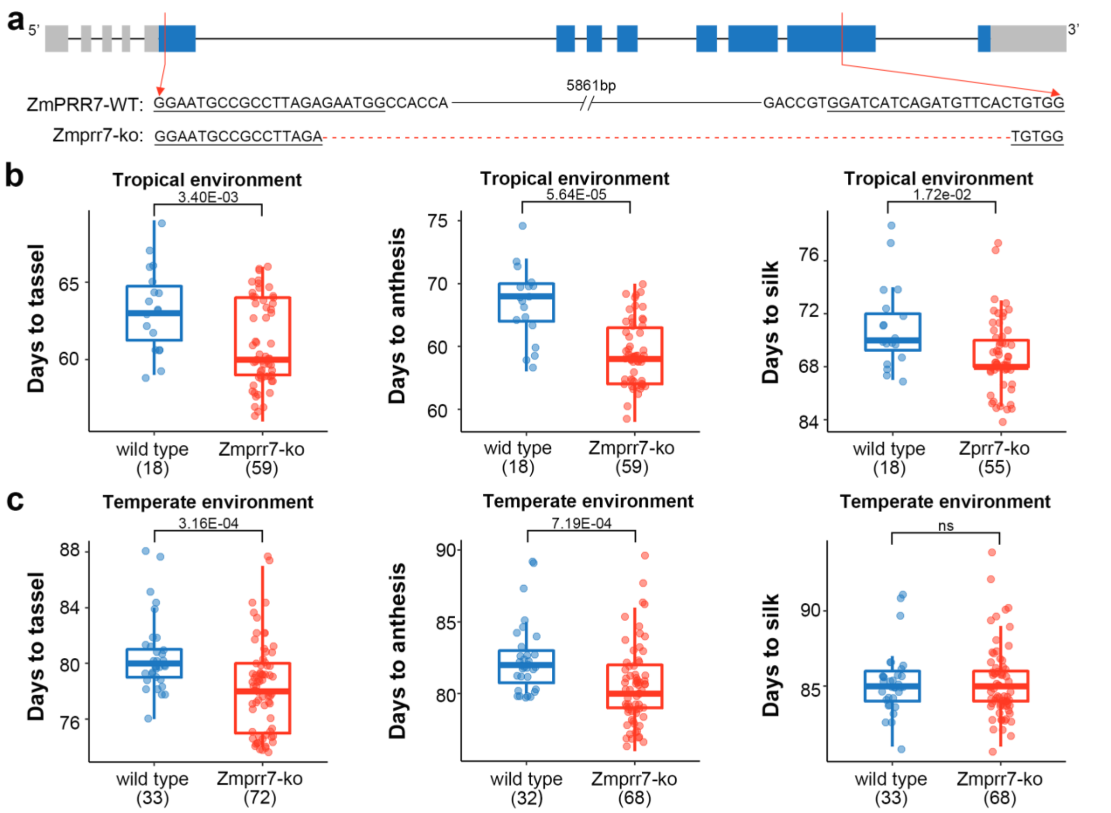


**Supplementary Figure 23. Phenotype analysis of CRISPR/Cas9 mutation for *ZmPRR7* in different environments. a,** Gene structure and sequences of *ZmPRR7* target regions in wild type, Zmprr7 CRISPR/Cas9 knockout mutants. **b,** Statics of days to tassel, days to anthesis, days to silk in Hainan province (2019; China; E109°, N18°; tropical environment). **c,** Statics of days to tassel, days to anthesis, days to silk in Jilin province (2020; China; E125°, N44°; temperate environment).

**Supplementary Table 8. Large inversions in *parviglumis* and *mexicana*.**

| Chromosome | Start | End | Population |
|---|---|---|---|
| 1 | 80,785,790 | 113,131,800 | *parviglumis* |
| 8 | 44,757,669 | 57,270,984 | *parviglumis* |
| 1 | 162,469,493 | 172,463,072 | *mexicana* |
| 2 | 183,082,698 | 197,102,429 | *mexicana* |
| 4 | 52,168,193 | 68,172,923 | *mexicana* |
| 4 | 170,740,777 | 194,384,324 | *mexicana* |
| 5 | 81,856,817 | 90,827,280 | *mexicana* |
| 5 | 104,117,103 | 114,775,228 | *mexicana* |
| 8 | 527,009 | 31,323,652 | *mexicana* |
| 8 | 61,141,607 | 75,742,589 | *mexicana* |
| 9 | 29,532,860 | 37,357,290 | *mexicana* |
| 9 | 120,347,800 | 144,538,800 | *mexicana* |
| 10 | 29,212,760 | 34,456,282 | *mexicana* |
| 10 | 79,158,738 | 94,363,919 | *mexicana* |

**Supplementary Table 23. Primers used in transgenic study.**

| Primer name | Primer sequence (5'-3') | Purpose |
|---|---|---|
| wox11-F | CTCCGAGCACAACGGCA | Detect *ZmWOX11* expression level |
| wox11-R | CCGAAAATGGCTCGCATGTC | Detect *ZmWOX11* expression level |
| J11-1F | GGCTACCTTGCTGCTCCTATC | Clone *ZmCOL9* CRISPR/Cas9 target region |
| J11-1R | CGCCCATGAGGAGAATTGGT | Clone *ZmCOL9* CRISPR/Cas9 target region |
| J11-2F | CTTTGCTGCTGGGCCATTTG | Clone *ZmCOL9* CRISPR/Cas9 target region |
| J11-2R | TGATAGGAGCAGCAAGGTAGC | Clone *ZmCOL9* CRISPR/Cas9 target region |
| J4-1F | CGTGGACTCCCTCTAGTTGC | Clone *ZmPRR7* CRISPR/Cas9 target region |
| J4-2R | CGCTACTTCCATTCTGCCCA | Clone *ZmPRR7* CRISPR/Cas9 target region |
